# Anxa1+ dopamine neuron vulnerability defines prodromal Parkinson’s disease bradykinesia and procedural motor learning impairment

**DOI:** 10.1101/2024.12.22.629963

**Authors:** Ioannis Mantas, Alessandro Contestabile, Vasiliki Skara, Camille Loiseau, Ines A Santos, Kaitlyn M L Cramb, Roberta Filograna, Richard Wade-Martins, Peter Magill, Konstantinos Meletis

**Affiliations:** Department of Neuroscience, Karolinska Institutet, Stockholm, Sweden; Medical Research Council Brain Network Dynamics Unit, Nuffield Department of Clinical Neurosciences, University of Oxford, Oxford OX1 3TH, United Kingdom; Oxford Parkinson’s Disease Centre, University of Oxford, Oxford OX1 3QX, United Kingdom; Oxford Parkinson’s Disease Centre and Department of Physiology, Anatomy and Genetics, University of Oxford, Dorothy Crowfoot Hodgkin Building, South Parks Road, Oxford OX1 3QU, UK; Kavli Institute for Nanoscience Discovery, University of Oxford, Dorothy Crowfoot Hodgkin Building, South Parks Road, Oxford OX1 3QU, UK; Department of Medical Biochemistry and Biophysics, Karolinska Institutet, Stockholm, Sweden; Aligning Science Across Parkinson’s (ASAP) Collaborative Research Network, Chevy Chase, MD 20815, United States of America

## Abstract

Progressive degeneration of dopamine neurons (DANs) defines Parkinson’s disease (PD). However, the identity and function of the most vulnerable DAN populations in prodromal PD remain undefined. Here, we identify substantia nigra DANs with Annexin A1 (Anxa1) expression as selectively vulnerable across multiple prodromal PD models and significantly reduced in patient-derived DANs. We found that Anxa1+ DANs have a unique functional profile, as they do not signal reward or reinforce actions, and they are not necessary for motivated behavior. Instead, activity of Anxa1+ DAN axons correlates with vigorous movements during self-paced exploration, yet their silencing only disrupts a subset of action sequences that mirror a PD bradykinesia profile. Importantly, Anxa1+ DANs are essential for procedural learning in a maze task and for motor learning of dexterous actions. These findings establish the early vulnerability of Anxa1+ DANs in PD, whose function can explain prodromal bradykinesia and impairments in procedural motor learning.

## Introduction

Parkinson’s disease (PD) is diagnosed based on clinical criteria, requiring the presence of bradykinesia along with at least one additional motor symptom, such as tremor, rigidity, or postural instability^1^. At the time of diagnosis, it is estimated that 60–70% of dopamine neurons (DANs) in the substantia nigra pars compacta (SNc) have already degenerated^2.^. The loss of SNc DANs is strongly associated with deficits in movement initiation and vigor, which underlie the characteristic motor impairments of PD^3^. Furthermore, the extent of SNc DAN degeneration correlates with the severity of bradykinesia and rigidity^4,5^. Prior to meeting diagnostic criteria, the prodromal phase of PD may include mild bradykinesia alongside emotional and cognitive impairments^6^, such as deficits in attention, visuospatial processing, memory, and executive function^7^. Anatomical studies have found that DANs in substantia nigra pars compacta (SNc) show earlier degeneration compared to DANs in the ventral tegmental area (VTA) in PD patients^8^. Progressive neurodegeneration in PD supports a differential vulnerability among DANs, beyond their localization to the VTA versus SNc, where the ventral tier of the SNc is particularly susceptible to degeneration^9^. Since PD patients are diagnosed after significant DAN degeneration has already occurred, our understanding of how the progressive DAN loss contributes to different stages of the disease remains limited.

Neuroanatomical and molecular classification of DANs using genetic strategies and single-cell RNA-seq methods has revealed an unexpected diversity^10–14^. A DA population defined by expression of the markers Sox6 and Aldh1a1 shows degeneration in PD patients^15–17^ and PD animal models^18–20^. This vulnerable population has a spatial organization corresponding to the SNc ventral tier domain and possibly overlaps with a vulnerable AGTR1+ DA subpopulation^21^. Interestingly, recent evidence suggests that this Sox6+/Aldh1a1+ subcluster within the SNc can be further subdivided into distinct subtypes^18^. However, the differential vulnerability of these subtypes throughout the degeneration process remains poorly understood. This highlights the need for high-resolution profiling of vulnerability-related transcriptional signatures to map specific DAN subtype degeneration trajectories, offering insights into their role in prodromal deficits.

Midbrain DANs show signals that can encode reward prediction errors^22,23^. In the reinforcement learning model, midbrain DANs function as the critic, broadcasting a scalar signal to the striatum that encodes the predicted value of actions^24,25^. However, growing evidence suggests that DANs do not function as an entirely homogeneous population that only signals reward prediction errors or value^26,27^; instead, their discrete neuroanatomical organization^13,14^ could reflect distinct roles in for example motor programs and learning^14,28–31^. For instance, VTA DANs projecting to nucleus accumbens (ACB) can signal motivational effort^32^, while SNc DANs projecting to the dorsal striatum (caudoputamen; CP), have been linked to movement initiation, invigoration and motor skill acquisition^33–35^. Adding complexity to the DA signals, for example in goal-directed motor programs DA signals can show ramping in ACB^36,37^, while DA signals can show wave-like patterns across dorsal CP subregions with learning^38^. Recent findings further indicate that action-potential-induced DA release facilitates reward-oriented behavior but is not essential for movement initiation^39,40^, further supporting functional specialization within the DA system. In summary, it remains unclear whether the functionally and topographically distinct DA signals are segregated in molecularly defined DAN subtypes and how their dysfunction could explain specific motor and non-motor deficits in PD models.

In this study, we used multiple PD mouse models that recapitulate progressive DAN degeneration as found in the prodromal stages of the disease, to define DAN subpopulations with early vulnerability. We describe here a molecularly and neuroanatomically specialized DAN subpopulation that degenerates early in several PD mouse models, is reduced in patient-derived DANs, and that is defined by expression of the marker Annexin a1 (Anxa1). We have genetically targeted and manipulated Anxa1-expressing DANs and establish their central role in procedural motor learning.

## Results

### Anxa1 is a marker of an early PD vulnerable SNc DAN subpopulation

To identify vulnerable DAN subtypes that degenerate in the initial stage of PD we employed multiple mouse models with mild or progressive DAN degeneration. First, we aimed to understand how DAN subtype markers changed in the early degeneration stage in two different models of DAN-specific mitochondrial deficiency: conditional Mfn2 KO mice (cMfn2KO^41^) and conditional Tfam mice (MitoPark^42^) (**Figure 1A**). For this reason, we performed RNA sequencing on FACS-sorted midbrain DANs from cMfn2KO mice three weeks post-recombination (cMfn2KO-early) and 10-12-week-old MitoPark (MP-early) mice, along with their respective controls (**Figure 1A**). Differential expression analysis of genes enriched in midbrain DAN subtypes revealed that both PD models showed a shared profile of significant early-stage downregulation of the Annexin A1 (Anxa1) transcript. (**Figure 1B**; **Figure S1**).

**Figure 1.**
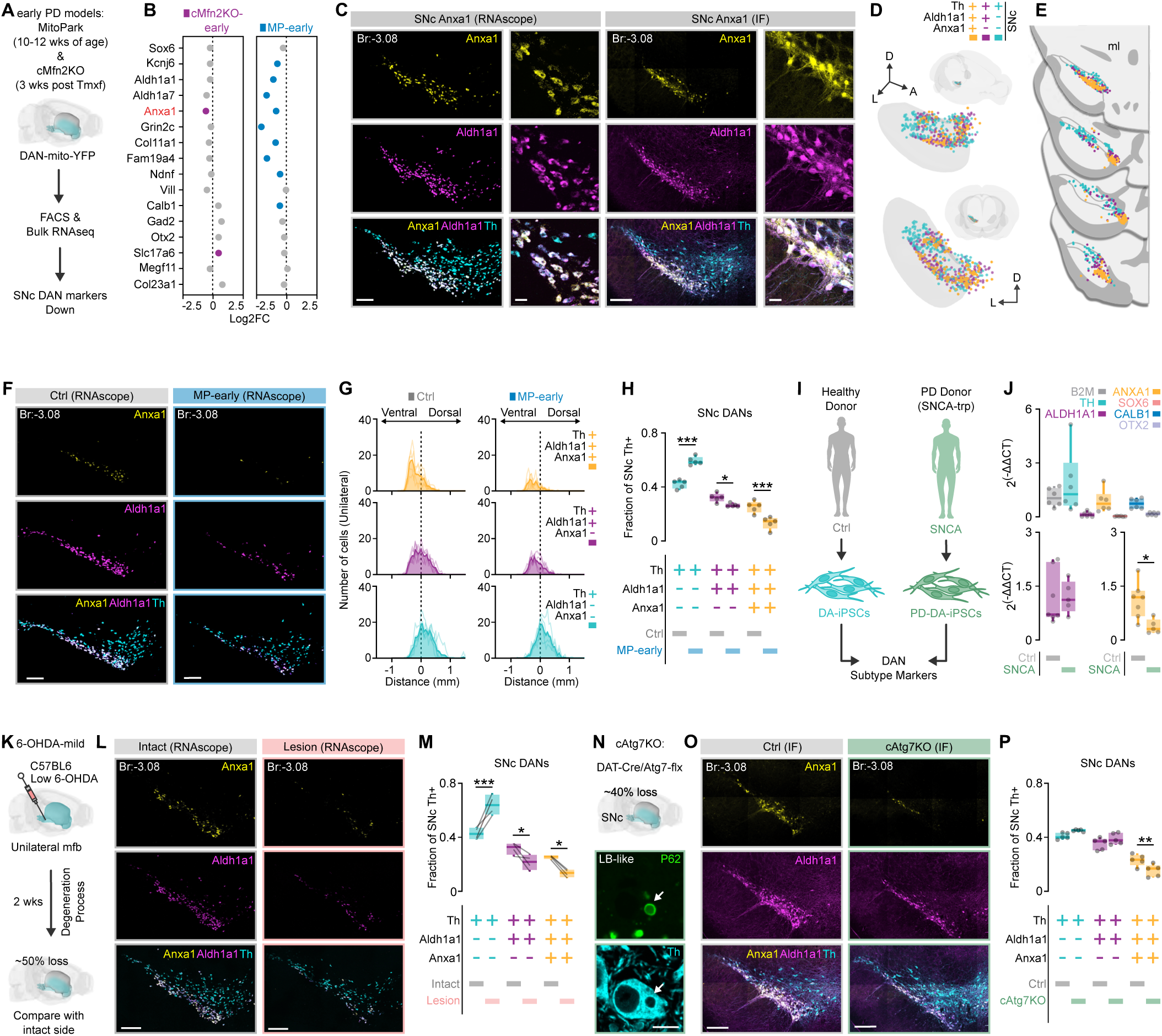
Anxa1 expression identifies an early PD-vulnerable subpopulation of SNc- DANs. **(A)** Schematic representation of FACS followed by bulk RNA-seq analysis of DANs from 10– 12-week-old MP-early (n = 5) and cMfn2KO-early (n = 5) mice, 3 weeks post-tamoxifen injection. **(B)** Graph displaying significantly downregulated DAN subtype markers (MP-early: blue; cMfn2KO-early: magenta; FDR < 0.05). **(C)** ISH (left) and IF (right) demonstrating co-expression of Anxa1, Aldh1a1, and Th in the SNc. Low- and high-magnification scale bars are 200 μm and 30 μm, respectively. **(D)** 3D reconstruction showing different DAN subpopulations visualized using anti-Anxa1, anti-Aldh1a1, and anti-Th antibodies in the SNc. **(E)** Coronal sections illustrating the spatial distribution of these subpopulations in the SNc. **(F)** ISH depicting Anxa1, Aldh1a1, and Th expression in the SNc of control (Ctrl) and MP-early mice. **(G)** Line graph showing the dorsoventral distribution of the three SNc DAN subpopulations in Ctrl (n = 5) and MP-early (n = 5) mice. **(H)** Boxplot illustrating the proportions of these subpopulations in Ctrl and MP-early mice (two-way ANOVA, Genotype × Subtype: F(2,24) = 49.66, p < 0.0001; *p = 0.0452, ***p < 0.0001, Sidak’s post-hoc test). **(I)** Illustration showing ANXA1 mRNA quantification in iPSC-DANs from PD patients carrying the SNCA-trp mutation. **(J)** Upper: Boxplots showing the expression of various SNc DAN subtype markers in iPSC- DANs. Lower: Boxplots comparing ALDH1A1 (left) and ANXA1 (right) mRNA levels between healthy controls and SNCA-trp iPSC-DANs. **(K)** Schematic representation of the low-concentration 6-OHDA lesion model. **(L)** ISH showing Anxa1, Aldh1a1, and Th expression in the SNc of intact and lesioned hemispheres. **(M)** Boxplot comparing the proportions of the three SNc DAN subpopulations in the intact versus lesioned hemispheres (n = 4; RM two-way ANOVA, Side × Subtype: F(2,9) = 36.33, p < 0.0001; *p < 0.05, ***p = 0.0002, Sidak’s post-hoc test). **(N)** Schematic illustration of cAtg7KO mice and IF images showing LB-like p62+ inclusions (arrows) in SNc DANs. **(O)** IF images showing the expression of Anxa1, Aldh1a1, and Th in the SNc of Ctrl and cAtg7KO mice. **(P)** Boxplot showing the proportions of the three SNc DAN subpopulations in Ctrl (n = 5) and cAtg7KO (n = 5) mice (two-way ANOVA, Genotype × Subtype: F(2,24) = 8.470, p = 0.0016; **p = 0.0082, Sidak’s post-hoc test). Boxplots display all data points, with the 25th and 75th percentiles (boxes), median (center), and maxima (whiskers). Abbreviations: PD: Parkinson’s Disease, SNc: Substantia Nigra pars compacta, DANs: Dopaminergic Neurons, FACS: Fluorescence-Activated Cell Sorting, RNA-seq: RNA Sequencing, Tmxf: tamoxifen, MP: MitoPark, cMfn2KO: Conditional Mitofusin 2 Knockout, FDR: False Discovery Rate, FC: fold change, Br: bregma, A: anterior, D: dorsal, L: lateral, ISH: In Situ Hybridization, IF: Immunofluorescence, ml: medial lemniscus, Ctrl: Control, iPSC- DANs: Induced Pluripotent Stem Cell-Derived Dopaminergic Neurons, CT: threshold cycle, SNCA-trp: SNCA-triplication, 6-OHDA: 6-Hydroxydopamine, mfb: medial forebrain bundle, cAtg7KO: Conditional Atg7 Knockout, LB: Lewy Bodies. See also **Figure S1**.

To gain a better understanding of the anatomical organization of Anxa1+ DANs, we mapped their distribution in relation to the vulnerability marker Aldh1a1^16^ (**Figure 1C**). We found that ∼99% of Anxa1+ DANs co-expressed Aldh1a1, while ∼40% of the Aldh1a1+ DANs expressed Anxa1 (**Figure 1C**; **Figure S1**). Using the medial lemniscus to define the SNc-VTA border, we found that the Anxa1+ DANs were localized in the ventral part of the SNc, and a small fraction was found in the VTA region (**Figure 1C-1E; Figure S1**). Moreover, we observed that Anxa1+ DANs were localized laterally in the anterior portions of the SNc, with their distribution becoming progressively more medial towards more posterior coronal planes (**Figure 1C-1E**).

To gain a more detailed understanding of the vulnerability of Anxa1+ DANs, we assessed Anxa1 expression in the SNc during early stages of degeneration and compared it to the expression of the putative vulnerability marker Aldh1a1^16^. To this end, we collected midbrain sections from MP- early mice and mapped the tissue expression of Anxa1 and Aldh1a1 in DANs. These experiments revealed a significant reduction in the number of Anxa1+ DANs in the ventral tier of the SNc in MP-early mice (**Figure 1F-1G**). Furthermore, we observed a significant decrease in the fraction of Anxa1+ DANs relative to the total Th+ cells in the SNc, whereas no reduction was found in the Anxa1+ DANs of the VTA (**Figure 1H, Figure S1**).

Building on our findings that Anxa1 serves as a vulnerability marker in mouse models, we further investigated the expression of ANXA1 in cells derived from PD patients. To do this, we used iPSC-derived DANs from PD patients with the SNCA triplication (SNCA-trp) mutation and from healthy donors (**Figure 1I**). First, we confirmed the expression of several genes that define different midbrain DAN subtypes, including ALDH1A1, CALB1, OTX2, and ANXA1 (**Figure 1J**). While ALDH1A1 expression was not significantly altered, we observed a significant reduction in ANXA1 expression in iPSC-DANs carrying the SNCA-trp mutation, further strengthening the link between ANXA1 downregulation and PD (**Figure 1J**).

To further investigate whether Anxa1 defines a vulnerable DAN subpopulation during the initial stages of PD as a common principle, we analyzed the degeneration of Anxa1+ DANs in two additional early-PD mouse models. Anxa1+ DAN degeneration in MitoPark and cMfn2KO mice, driven by mitochondrial deficits, prompted us to explore two other mild PD models: a mild toxin model and a mild progressive autophagy deficiency model. To capture vulnerability differences across the entire midbrain DAN population, we used a mild toxin model based on a low dose 6- hydroxydopamine (6-OHDA) injection into the medial forebrain bundle. This approach established a mild degeneration model, resulting in partial loss (∼50%) of Th+ DANs (**Figure S1**). In this model, we observed a significant decrease in the Anxa1+ fraction in the SNc compared to the intact side (**Figure 1K–1M**). To study progressive degeneration associated with dopamine-specific autophagy deficiency, we analyzed 24-week-old conditional Atg7 knockout mice (cAtg7KO) with Lewy body-like inclusions^43^ (**Figure 1N**). In the SNc of cAtg7KO mice, we observed a specific and significant reduction in Anxa1+ DANs (**Figure 1O and 1P**). Notably, analyses of the VTA in both models revealed no significant effects on the Anxa1+ fraction (**Figure S1**). Together, these findings demonstrate that SNc Anxa1+ DANs represent a particularly vulnerable subpopulation, exhibiting early degeneration across multiple PD models regardless of the underlying mechanism.

### Specialized organization of Anxa1+ DANs

To elucidate the organization and function of Anxa1+ DANs, we generated a mouse line with specific expression of Flp recombinase (Flp) in Anxa1-expressing cells (Anxa1-flp mice) (Figure 2A). We injected a Flp-dependent AAV to express the marker eYFP (AAV fDIO-eYFP) into the SNc of Anxa1-flp mice, enabling genetic labeling of Anxa1+ DANs (**Figure 2A**–**2B**; **Figure S2**). First, we mapped the dendritic processes of Anxa1+ DANs within the Substantia nigra pars reticulata (SNr) (**Figure 2C**–**2E**). We observed that Anxa1+ DANs extend long dendrites deep into the anterior and medial portions of the SNr (**Figure 2C**–**2E**). Examining long-range projections to the striatum, we found that Anxa1+ DANs formed a dense projection to the most dorsal domain of the CP (dCP) and some axonal labeling in the core and medial shell of the ACB (**Figure 2F**–**2H**). Anxa1+ axonal labeling revealed a pronounced density gradient enriched in the anteromedial regions of the dCP (**Figure 2F**–**2H**), aligning with Anxa1 immunoreactivity observed in the dCP (**Figure S2**). The Anxa1+ projection was concentrated in the most dorsal region of the broader Aldh1a1 immunoreactive territory (**Figure S2**), with the projection pathway traversing the dorsolateral region of the lateral hypothalamic area, corresponding to the location of the nigrostriatal component of the medial forebrain bundle (**Figure S2**). The spatial organization of Anxa1+ DANs and their discrete projection patterns suggest a specialized role in specific behaviors mediated by basal ganglia (BG) loops involving the most dorsal domains across the dorsomedial and dorsolateral striatum.

**Figure 2.**
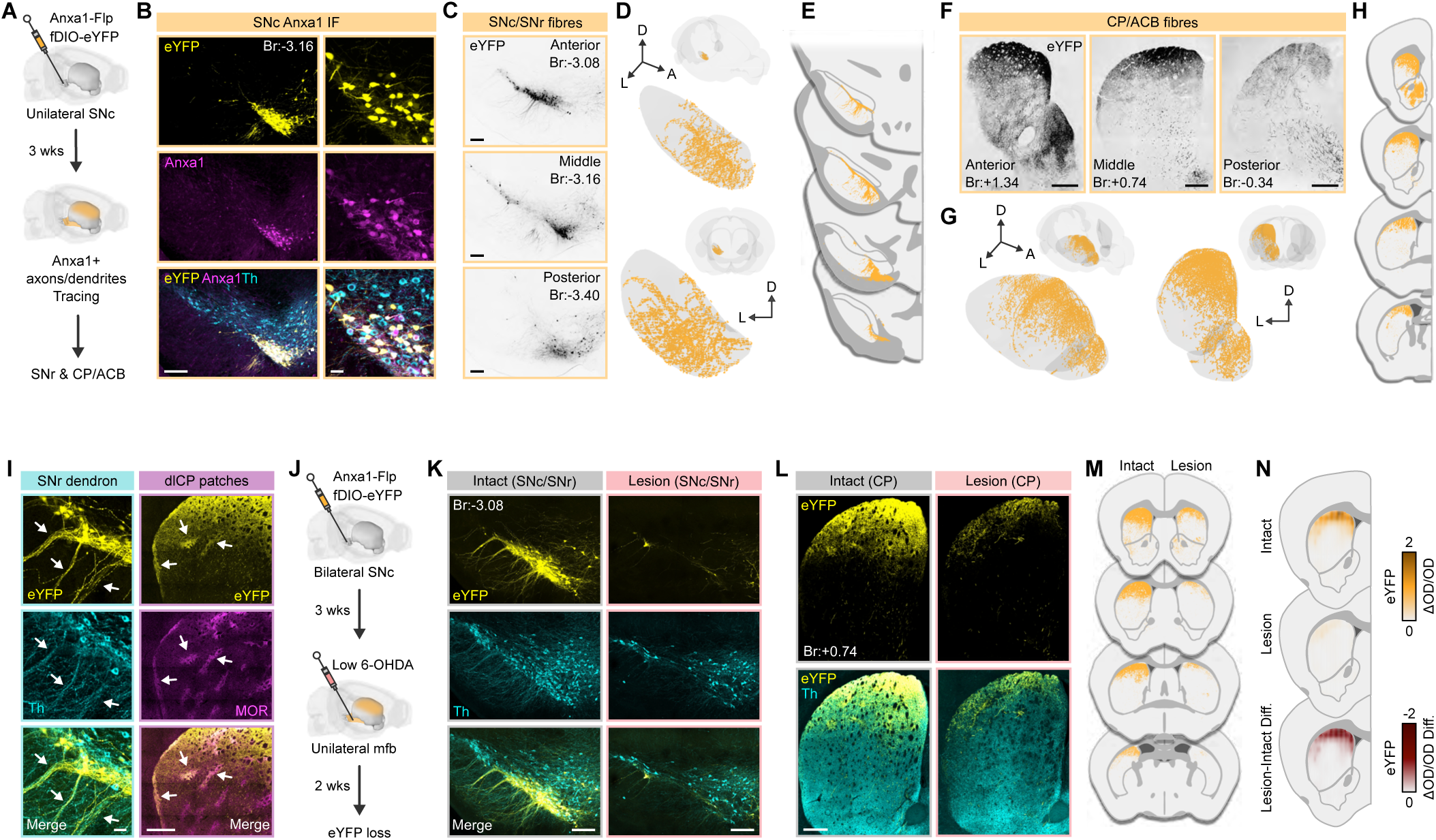
The specialized organization of Anxa1+ DANs and their projections to a dorsal domain in CP. **(A)** Schematic illustration representing the labeling of Anxa1+ fibers in Anxa1-Flp mice. **(B)** IF images showing the overlap between eYFP labeling and Anxa1-positive dopaminergic neurons (Anxa1+ DANs). Low-magnification scale bar: 200 μm; high-magnification scale bars: 30 μm. **(C)** Fluorescent images of eYFP+ labeled fibers and somata at various anteroposterior coronal levels within the SNc and SNr. **(D)** 3D reconstruction of eYFP+ labeled fibers localized in the SNr. **(E)** Schematic coronal sections illustrating the distribution of eYFP+ labeled fibers in the SNr. **(F)** Fluorescent images displaying eYFP+ labeled axons at distinct anteroposterior coronal levels within the CP and ACB. **(G)** 3D reconstruction of eYFP+ labeled fibers within the CP and ACB. **(H)** Schematic coronal sections showing the distribution of eYFP+ labeled fibers within the CP and ACB. **(I)** IF images illustrating eYFP+ labeled fibers in dendron bouquets within the SNr (left; scale bar: 30 μm) and in dorsolateral CP patches and subcallosal streak (right; scale bar: 500 μm). **(J)** Schematic diagram depicting bilateral Anxa1+ fiber labeling in combination with low-concentration 6-OHDA treatment. **(K)** IF images showing eYFP labeling alongside Th immunoreactivity in the SNc of the intact (left) and lesioned (right) sides (scale bars: 200 μm). **(L)** IF images depicting eYFP labeling alongside Th immunoreactivity in the CP of the intact (left) and lesioned (right) sides (scale bars: 500 μm). **(M)** Schematic coronal sections showing the distribution of eYFP+ labeled fibers within the CP and ACB of intact and lesioned side. **(N)** Heatmap showing the fluorescent intensity of eYFP in the intact and lesioned sides, with the difference in intensity displayed (n = 5 mice). Abbreviations: eYFP: enhanced Yellow Fluorescent Protein, DANs: Dopaminergic Neurons, SNc: Substantia Nigra pars compacta, SNr: Substantia Nigra pars reticulata, CP: Caudoputamen, A: anterior, D: dorsal, L: lateral, dlCP: dorso-lateral CP, ACB: Accumbens, Br: bregma, 6-OHDA: 6- Hydroxydopamine, mfb: medial forebrain bundle, OD: optic density, Diff.: difference. See also **Figure S2**.

To investigate the DA release profile in the dCP, a domain characterized by dense Anxa1+ projections, we compared it to a more central domain (cCP) with sparse Anxa1+ projections. DA recordings were performed in the dCP and cCP using a genetically encoded DA sensor (dLight1.3b^44^) combined with optogenetic stimulation of Anxa1+ DANs via Flp-dependent ChRmine^45^ expression (**Figure S2**). Optogenetic activation of Anxa1+ DANs at varying intensities elicited DA release in the dCP (**Figure S2**). Notably, optogenetically evoked DA release in the cCP exhibited a lower amplitude compared to the dCP signal (**Figure S2**), highlighting the specialized role of Anxa1+ DANs in shaping DA dynamics within specific CP subregions.

A detailed analysis of eYFP labeling revealed that the Anxa1+ DAN axons and dendrites exhibit features characteristic of the striosomal network (**Figure 2I**). Specifically, we observed that Anxa1+ dendrites in the anterior SNr had a dendron-like structure^46^ (**Figure 2I**), and that eYFP axonal labeling in the dorsolateral CP was enriched in the mu-opioid receptor (MOR) immunoreactive striosomes and subcallosal streak, suggesting a specialized nigro-striato-nigral loop organization (**Figure 2I**).

To directly assess the vulnerability and degeneration of Anxa1+ DANs and their axons, we combined genetic labeling using bilateral SNc injections of Flp-dependent eYFP in Anxa1-Flp mice with a low-dose unilateral 6-OHDA injection into the medial forebrain bundle (**Figure 2J**). Degeneration of the eYFP+ SNc neurons was accompanied by near-complete ablation of the corresponding eYFP+ axons in the dCP across the anteroposterior axis (**Figure 2K**–**2N**). Together, our findings establish that Anxa1+ DANs form a specialized circuit that defines PD-vulnerable territories in CP.

### Anxa1+ DANs do not encode motivational value

To evaluate the response of Anxa1+ axons to natural rewards, we recorded calcium transients from Anxa1+ projections in the dCP using the genetically encoded calcium sensor GCaMP8m (**Figure 3A and** **3B**). In addition, we recorded calcium transients from general DAN projections (i.e. using DAT-cre mice) in the dCP and measured DA release using the dLight1.3b sensor in both dCP and cCP (**Figure 3A and** **3B**). To capture reward-related responses, we used water-deprived mice provided with access to a licking spout (**Figure 3C; Supplementary Video1**). The recorded signals were synchronized with licking behavior, tracked via a high-speed camera, to detect reward-related activity in a freely moving setup (**Figure 3C and** **3D**). Recordings showed that the Anxa1+ as well as the DAT+ axons in dCP exhibited a ramping activity that peaked upon contact with the water droplet, followed by a rapid return to baseline during water consumption (**Figure 3D**-**3G**). Similarly, the recorded DA signals in the dCP showed the same dynamics (**Figure 3E**-**3G**). In contrast, recordings from the cCP revealed a typical reward-related phasic DA release profile, showing a ramping signal before droplet contact with a peak during reward consumption that then returned to baseline (**Figure 3E**-**3G**). This suggests that the activity of Anxa1+ projections to CP does not correlate with the classical value or reward prediction signals, indicating a distinct role in other DA-related processes.

**Figure 3.**
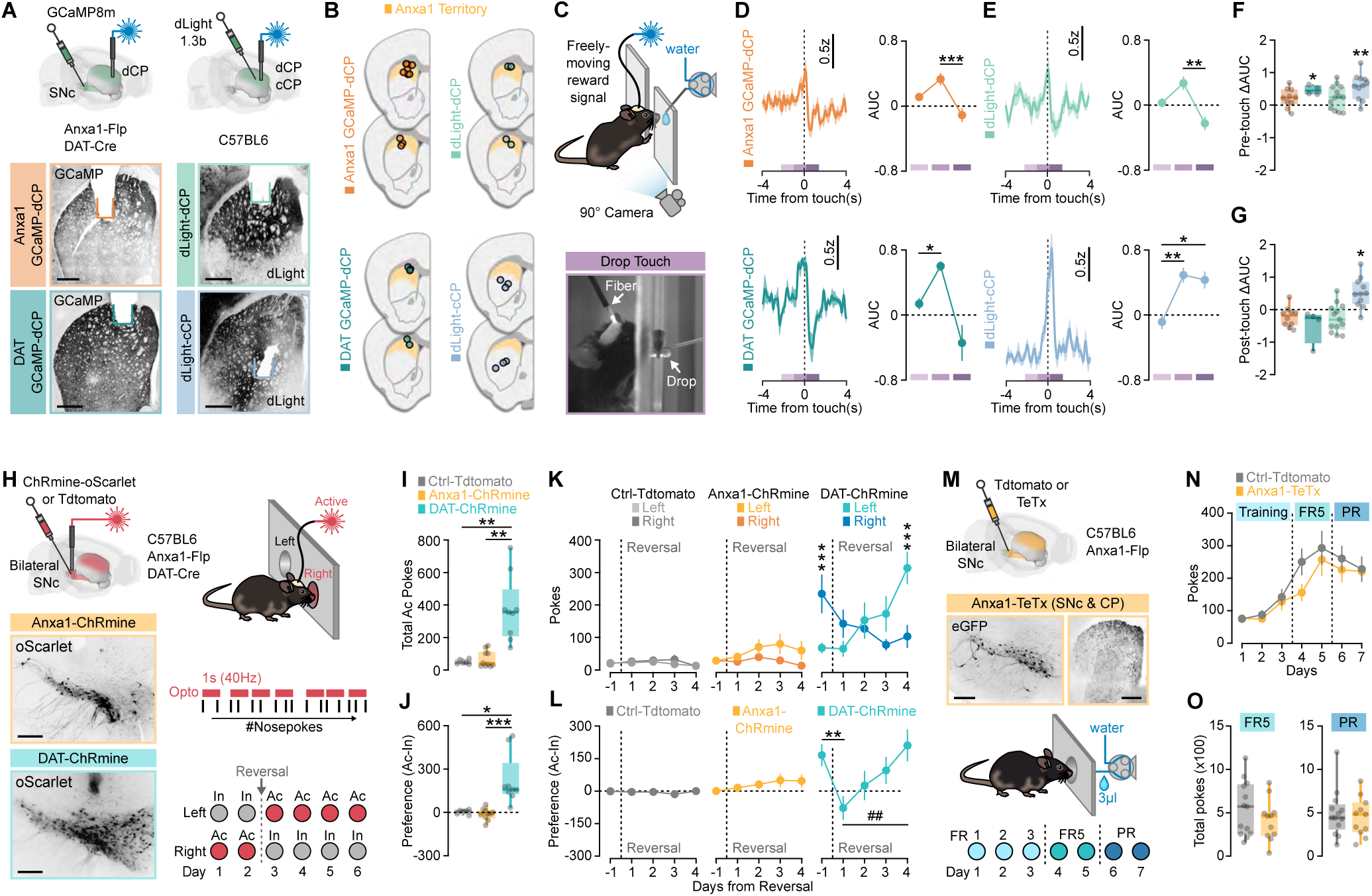
Anxa1+ DANs do not encode motivational value. **(A)** Illustration (top) and fluorescent images (bottom) depicting the various groups of mice utilized for FP recordings in the CP (scale bars: 500 μm). **(B)** Schematic representation of coronal sections illustrating the recording locations within the CP for each experimental group. **(C)** Illustration (top) and image (bottom) demonstrating the experimental setup for FP recordings during reward delivery. **(D)** Z-scores of FP signals aligned to the water drop touch across all recordings from the Anxa1 GCaMP-dCP (mice = 7, n = 10 recordings) and DAT GCaMP-dCP (mice = 3, n = 4 recordings) groups. Line graphs represent average AUC across three time windows relative to the water drop touch for Anxa1 GCaMP-dCP (RM one-way ANOVA, F(1.725, 15.52) = 13.26, p = 0.0006; second vs. third: ***p = 0.0003, Sidak’s post-hoc test) and DAT GCaMP-dCP (RM one-way ANOVA, F(1.093, 3.278) = 10.39, p = 0.0421; first vs. second: p = 0.0228, Sidak’s post-hoc test). **(E)** Z-scores of FP signals aligned to the water drop touch for recordings from the dLight-dCP (mice = 6, n = 13 recordings) and dLight-cCP (mice = 6, n = 11 recordings) groups. Line graphs illustrate average AUC across three time windows relative to water drop touch for dLight-dCP (RM one-way ANOVA, F(1.918, 23.01) = 9.658, p = 0.001; second vs. third: *p = 0.0037, Sidak’s post-hoc test) and dLight-cCP (RM one-way ANOVA, F(1.829, 18.29) = 10.58, p = 0.0011; first vs. second: *p = 0.002, first vs. third: p = 0.0228, Sidak’s post-hoc test). **(F-G)** Comparison of AUC differences for the second- and third-time windows relative to the first (baseline) across all recorded groups of mice (one-sample two-tailed t-test; second DAT GCaMP- dCP: p = 0.0308, second dLight-cCP: *p = 0.0028, third dLight-cCP: p = 0.0148, Bonferroni correction applied). **(H)** Left: Illustration (top) and fluorescent images (bottom) depicting ChRmine-oScarlet expression in the SNc of DAT-ChRmine and Anxa1-ChRmine mice (scale bar: 300 μm). Right: Illustration of the experimental setup and timeline for the self-stimulation protocol. **(I)** Boxplot comparing active pokes during the first two days of the self-stimulation protocol across groups (Ctrl-Tdtomato: n = 6, Anxa1-ChRmine: n = 10, DAT-ChRmine: n = 9; Kruskal-Wallis test, p = 0.0007; Ctrl-Tdtomato vs. DAT-ChRmine: *p = 0.004, Anxa1-ChRmine vs. DAT-ChRmine: *p = 0.0029, Dunn’s post-hoc test). **(J)** Boxplot comparing preference for the stimulation-associated poke during the first two days of the self-stimulation protocol across groups (Ctrl-Tdtomato: n = 6, Anxa1-ChRmine: n = 10, DAT- ChRmine: n = 9; Kruskal-Wallis test, p = 0.0003; Ctrl-Tdtomato vs. DAT-ChRmine: p = 0.0131, Anxa1-ChRmine vs. DAT-ChRmine: **p = 0.0003, Dunn’s post-hoc test). **(K)** Line graphs depicting the number of left and right pokes across pre- and post-reversal experimental days for Ctrl-Tdtomato, Anxa1-ChRmine, and DAT-ChRmine mice (RM two-way ANOVA, Side × Days: F(4, 32) = 15.70, p < 0.0001; Right vs. Left: **p < 0.001, Sidak’s post-hoc test). **(L)** Line graphs showing preference for the stimulation-associated port across pre- and post-reversal experimental days for Ctrl-Tdtomato, Anxa1-ChRmine, and DAT-ChRmine groups (Friedman test, p = 0.0004; day −1 vs. day 1: *p = 0.008 and day 1 vs. day 4: ##p = 0.0011, Dunn’s post-hoc test). **(M)** Illustration (top) and fluorescent images (middle) depicting TeTx-GFP expression in the SNc and CP of Anxa1-TeTx mice (scale bars: 300 μm and 500 μm). Bottom: Illustration demonstrating the experimental setup and timeline for the fixed ratio (FR) and progressive ratio (PR) protocol. **(N)** Line graphs presenting the number of pokes during the FR and PR protocols across experimental days for the two groups. **(O)** Boxplots comparing total pokes in the FR5 (left) and PR (right) stages of the protocol between the two groups. Data in line graphs are expressed as mean ± SEM. Boxplots depict all data points, the 25th and 75th percentiles (box), the median (center), and the maxima (whiskers). Abbreviations: DANs: Dopaminergic neurons, FP: Fiber photometry, dCP: dorsal Caudoputamen, cCP: central Caudoputamen, DAT: Dopamine Transporter, AUC: Area Under the Curve, SNc: Substantia Nigra pars compacta, Ctrl: Control, Opto: optogenetic stimulation, TeTx: Tetanus Toxin, eGFP: enhanced Green Fluorescent Protein, Ac: active, In: inactive, FR: Fixed Ratio, PR: Progressive Ratio. See also **Supplementary Video1**.

To better define the causal role of Anxa1+ DANs in value-guided or motivated behaviors, we employed two distinct perturbation approaches. We first used an optogenetic self-stimulation paradigm and compared the activation of Anxa1+ DANs with all DANs (DAT+ neurons). To target SNc DANs, we expressed the opsin ChRmine using cell-type specific expression in Anxa1-Flp or DAT-cre mice (**Figure 3H**). The self-stimulation protocol involved two 15-minute sessions with optogenetic activation during right nosepokes, followed by four sessions with activation during left nosepokes (**Figure 3H**). We found that optogenetic activation of DAT+ neurons strongly reinforced self-stimulation during the first two days of the task (**Figure 3I and 3J**). When active and inactive port assignment was reversed, DAT-Cre mice quickly reversed their responding from the inactive (right) to the active (left) port (**Figure 3K and 3L**). In contrast, optogenetic activation of the Anxa1+ DANs did not support self-stimulation behavior, and mice did not develop any significant preference for the active port during the last days of the reversal (**Figure 3I**-**3L**). To further determine the role of Anxa1+ DANs for motivational value, we used a fixed and progressive ratio schedule in mice with silenced Anxa1+ DAN neurotransmission using cell-type specific expression of the tetanus toxin light chain^47^ (**Figure 3M**). Mice with genetic silencing of Anxa1+ DANs exhibited no significant changes in the total number of nosepokes during FR5 or PR tasks, suggesting that Anxa1-silenced mice do not exhibit motivational deficits (**Figure 3N and** **3O**). Overall, our data show that Anxa1+ DANs do not shape reward-related or motivated behaviors and suggest that they instead contribute to other aspects of DA-related behaviors.

### Anxa1+ DAN projections are activated during vigorous actions

It has been previously described that calcium transients in DANs axons in dCP are associated with self-paced movement initiation and acceleration^30,33,48^. To test the role of Anxa1+ DANs in locomotion and motor programs, we recorded the activity of Anxa1+ axons in dCP during exploration of an open field (**Figure 4A**). We tracked locomotion patterns, linear speed and acceleration on a sub-second scale together with recording of calcium and DA transients. We found a significant positive correlation between Anxa1+ axonal activity in the dCP and linear speed, and this correlation was similar when we recorded DA release in the dCP (**Figure 4B**). The recorded DA release in cCP instead showed a negative correlation with locomotion speed (**Figure 4B**).

**Figure 4.**
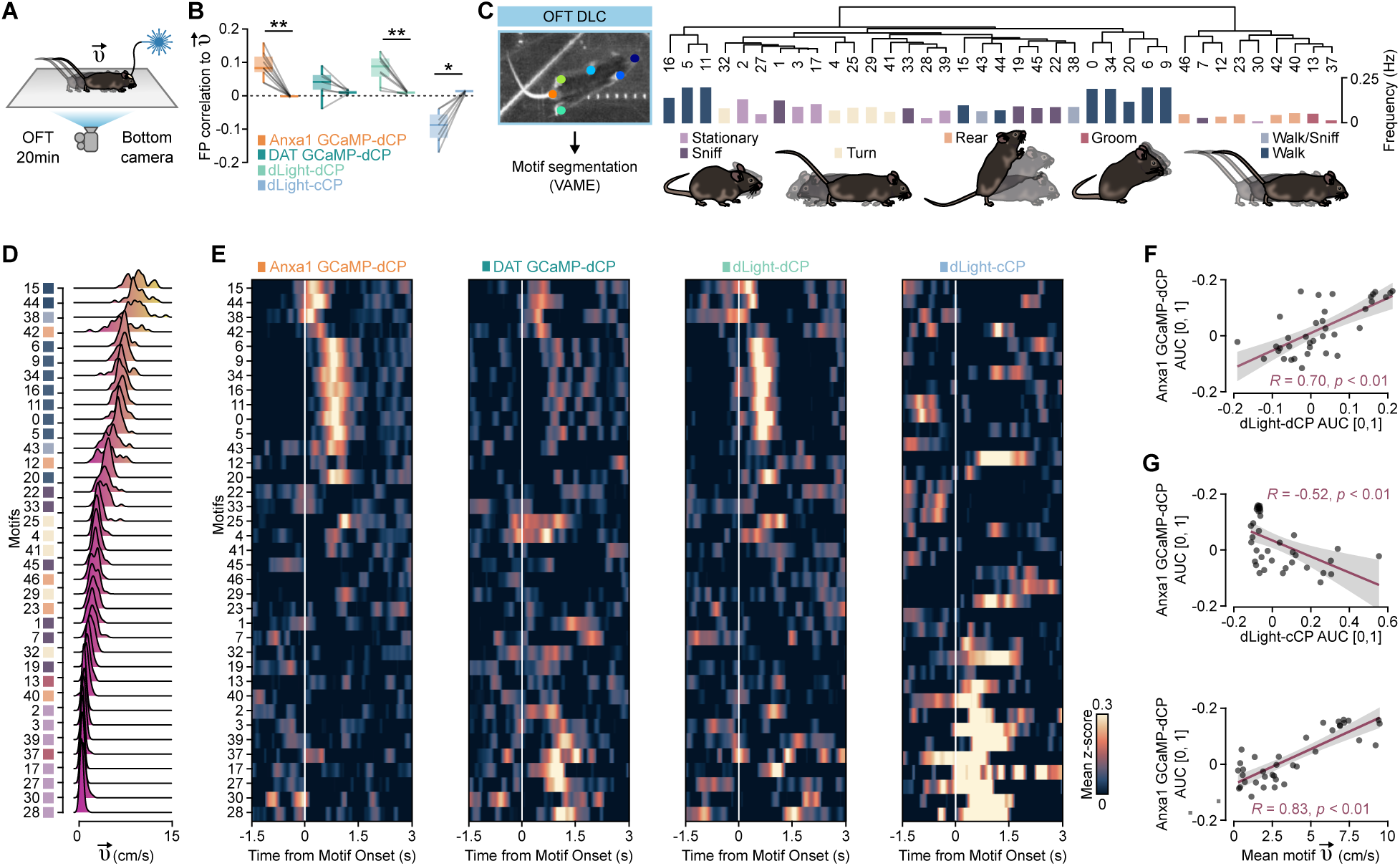
Anxa1+ DAN striatal axon activity tracks high-velocity behavioral motifs. **(A)** Illustration showing the experimental setup of the open field arena. Mouse behavior was recorded from below for 20 minutes while neuronal activity was measured using FP. **(B)** Boxplots comparing the correlation between the FP signal and linear speed (left boxplot) with the correlation between the shuffled signal and linear speed (right boxplot) for Anxa1 GCaMP- dCP (mice = 9, n = 9 recordings, **p = 0.0039, two-sided paired t-test), DAT GCaMP-dCP (mice = 5, n = 7 recordings), dLight-dCP (mice = 6, n = 8 recordings, **p = 0.0078, two-sided paired t- test), and dLight-cCP (mice = 6, n = 7 recordings, *p = 0.016, two-sided paired t-test). **(C)** Left: Image showing the pose estimation of six body parts using DLC. Right: Dendrogram depicting the hierarchical clustering of 37 behavioral motifs using VAME. Motifs are color-coded according to the simplified action categories: Stationary/Sniffing, Turn, Rear, Groom, Walk, and Sniff/Walk. **(D)** Graph showing the linear speed distribution for each behavioral motif. **(E)** Heatmaps displaying the mean z-scored FP signals (GCaMP and dLight) aligned to the onset of each motif. Motifs are ranked based on speed, as in d. **(F-G)** Graphs showing the correlation of FP signal AUC (time window: 0 to 1 s) between Anxa1-GCaMP-dCP and dLight-dCP (**F**, R = 0.70, p < 0.01) or dLight-cCP (**G**, R = −0.52, p < 0.01). **(H)** Graph showing the correlation between FP signal AUC (time window: 0 to 1 s) for Anxa1-GCaMP-dCP and mean motif linear speed (R = 0.83, p < 0.01). Boxplots show all data points, with the 25th and 75th percentiles (box), the median (center), and the maxima (whiskers). Abbreviations: OFT: open filed test, DANs: Dopaminergic neurons, FP: Fiber photometry, DAT: Dopamine transporter, dCP: dorsal Caudoputamen, cCP: central Caudoputamen, DLC: DeepLabCut, VAME: Variational Animal Motion Embedding, AUC: Area under the curve, R: Pearson correlation coefficient. See also **Figure S3** and **Supplementary Video2**.

To provide a more in-depth description of the correlation between Anxa1+ axon activity and kinematics, we used DeepLabCut (DLC)^49^ combined with an unsupervised probabilistic deep learning framework that identifies behavioral structure from deep variational embeddings of animal motion (VAME)^50^. This unsupervised analysis of postural dynamics identified 37 kinematic motifs, which we categorized into seven communities of actions: walk/sniff, walk, groom, turn, rear, sniff, and stationary (**Figure 4C**). To further understand the relationship between activity in Anxa1+ projections and locomotor speed, we ranked the motifs according to their respective speed distribution (**Figure 4D**) and aligned the calcium or DA signal with the onset of the corresponding motifs (**Figure 4E**). This analysis demonstrated that the activity of Anxa1+ projections was associated with high-speed motifs independently of the action type, and this profile was also observed in DA signals recorded in the dCP (**Figure 4E; Figure S3**). However, DA signals recorded in the cCP demonstrated a different pattern with strong activation after the onset of low-speed motifs (**Figure 4E; Figure S3**). Notably, the area under the curve (AUC, 0 to 1 sec) calculated for each motif from Anxa1-terminal activity showed a positive correlation with the AUC of dLight signals recorded in the dCP (**Figure 4F**) and a negative correlation with the AUC of dLight signals recorded in the cCP (**Figure 4G**). Furthermore, the AUC from Anxa1-terminal activity was highly positively correlated with motif speed (**Figure 4H**). These findings demonstrate a strong correlation between Anxa1+ projection activity and behaviors characterized by vigorous, high-velocity movements.

### Anxa1+ DAN silencing produces a prodromal PD bradykinetic profile

Given the strong correlation between Anxa1+ projection activity and high-velocity movements, we aimed to determine the causal role of Anxa1+ DANs in driving these motor actions and their relevance to prodromal motor deficits in mild or progressive PD mouse models. To investigate this, we genetically silenced Anxa1+ DANs using cell-type-specific expression of tetanus toxin light chain (**Figure 5A**) and compared with two disease stages: an early-stage PD (mild 6OHDA model and 10-12-week-old MitoPark mice) and a middle-stage PD (15-18-week-old MitoPark mice; **Figure 5A; Figure S4**). We first analyzed classical locomotor parameters during open field exploration, and we found that Anxa1-tetanus mice showed moderate reductions in linear and angular speed, accompanied by a slight increase in immobility time (**Figure 5B**-**5F**). The mild locomotor phenotype and acceleration distribution pattern in Axna1-silenced mice was similar to the early-stage PD models suggesting that loss of Anxa1+ DANs is linked primarily to a bradykinetic rather than an akinetic/rigid profile (**Figure 5E**-**5F**).

**Figure 5.**
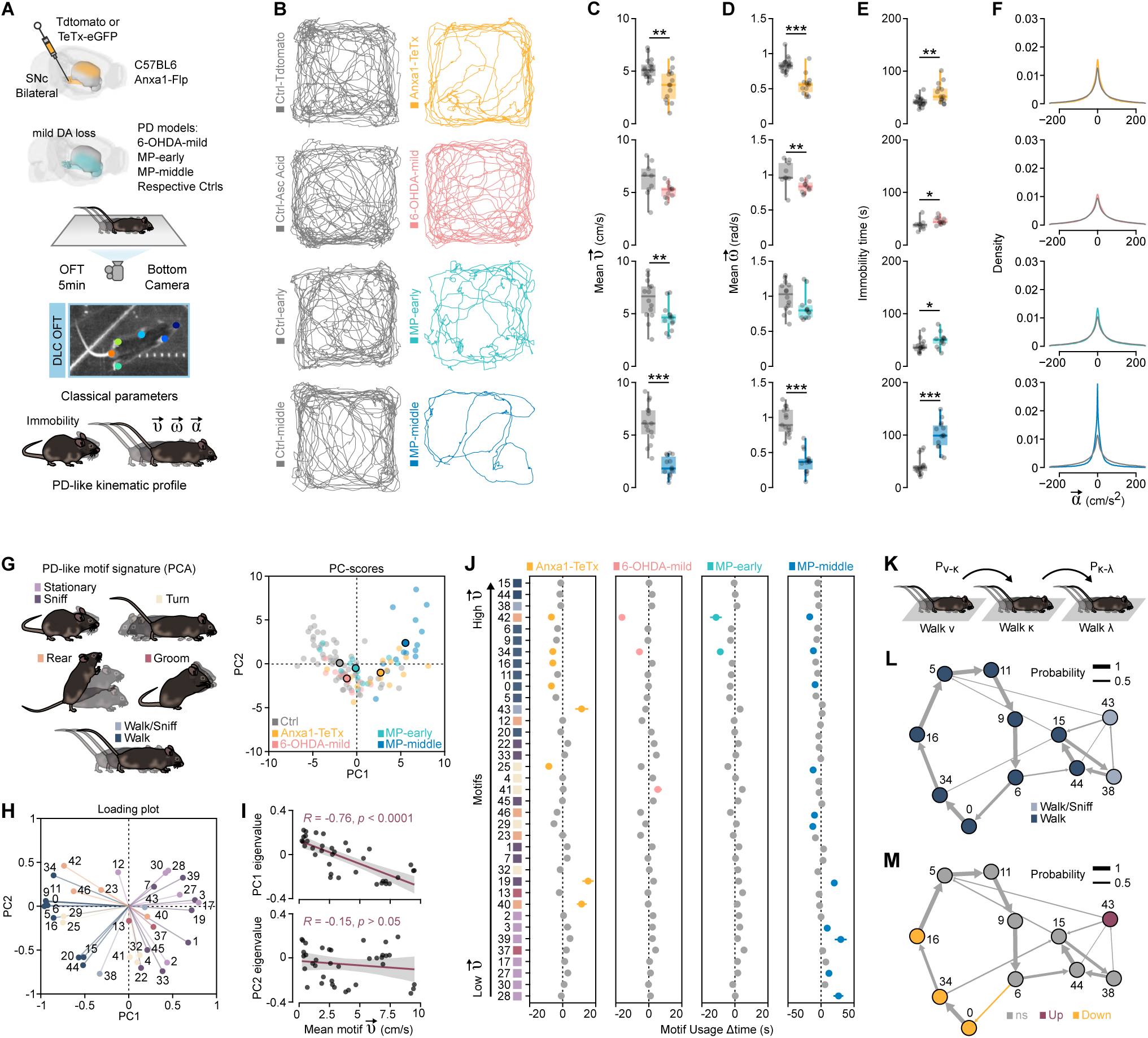
Genetic silencing of Anxa1+ DANs produces an early-stage PD bradykinesia profile. **(A)** Illustration depicting the distinct groups of mice, and the experimental setup utilized for the open field test. **(B)** Representative path-tracking traces of the various experimental groups during the open field test. **(C–E)** Boxplots presenting the mean linear speed **(C)**, mean angular speed **(D)**, and immobility time **(E)** for each control group (Ctrl-Tdtomato: n = 18, Ctrl-Asc Acid: n = 9, Ctrl-early: n = 14, Ctrl-middle: n = 15) and experimental group (Anxa1-TeTx: n = 14, 6OHDA-low: n = 11, MP-early: n = 13, MP-middle: n = 14). Statistical significance is indicated as *p < 0.05, **p < 0.01, ***p < 0.001 (two-sided unpaired t-test). **(F)** Graphs showing the acceleration distribution density for each group of mice. **(G)** Left: Illustration categorizing mouse behavior into five action motifs using VAME: Stationary/Sniffing, Turn, Rear, Groom, and Walk & Sniff/Walk. Right: PCA of motif usage, reporting the PC1 and PC2 for each group of mice. **(H)** Loading plot displaying PC1 and PC2 values for the different behavioral motifs, color-coded according to the action categories defined in (G). **(I)** Graphs depicting linear regression analyses between the mean motif speed and the eigenvalues of PC1 (top; R = −0.762, p < 0.0001) or PC2 (bottom; R = −0.1494, p = 0.3772). **(J)** Graphs illustrating the motif usage (Δtime) across different groups (data expressed as mean ± SEM). Motifs are arranged from bottom to top according to increasing speed. Colored symbols represent statistically significant altered motifs (Anxa1-TeTx: yellow, 6-OHDA-low: pink, MP-early: light blue, MP-middle: dark blue; multiple two-sided unpaired t-tests, adjusted p < 0.05, Holm-Sidak method). **(K)** Illustration showing the probability of one “walk” motif occurring after the other. **(L)** Statemap depiction of “walk” motifs (as nodes) and transition probabilities (as weighted edges) from control mice. **(M)** Statemap depiction of “walk” motifs (as nodes) and transition probabilities (as weighted edges) from Anxa1-TeTx mice (n = 3 mice; red: significant upregulated transition, yellow: significant downregulated transition; Fisher’s Exact Test, adjusted p < 0.05, Benjamini-Hochberg correction). The colored nodes represent the motifs with statistically significant altered usage in Anxa1-TeTx mice according to **(J)** (red: significant upregulated motif, yellow: significant downregulated transition motif). Boxplots display all data points, with the 25th and 75th percentiles (box), the median (center), and the maxima (whiskers). Abbreviations: DANs: Dopaminergic neurons, PD: Parkinson’s disease, OFT: open field test, Ctrl: Control, Asc: ascorbic, TeTx: Tetanus Toxin, 6-OHDA: 6-hydroxydopamine, MP: MitoPark, DLC: DeepLabCut, PCA: Principal component analysis, PC: Principal component, R: Pearson correlation coefficient. See also **Figure S4** and **S5.**

Since the basic locomotor analysis did not allow us to understand in detail how Anxa1+ DANs can shape discrete motor programs, we decided to produce a detailed segmentation of the open field behavior and identify discrete and hierarchically organized action motifs (DLC^49^ and VAME^50^). Using this approach, we generated a kinematic signature for Anxa1-silenced mice and compared it with the behavioral structure of the mild and progressive PD models. To first identify the variables that could explain the kinematic differences across PD models and Anxa1-silenced mice, we performed a principal component analysis (PCA) using motif usage time. We found that the first principal component (PC1) effectively separated the groups according to PD stage (**Figure 5G**). The motif usage profile of Anxa1-tetanus mice positioned them between early and middle PD models along the PC1 axis (**Figure 5G**). Furthermore, we generated a loading plot to assess the influence of each motif usage on the principal components. “Walk” and “stationary” motifs were prominently represented at the opposite extremes of the PC1 axis (**Figure 5H**). Interestingly, linear regression analysis between motif speed and PC1 eigenvalues revealed a significant negative correlation (**Figure 5I**). In contrast, we did not observe any significant correlation between the eigenvalues of PC2 (**Figure 5I**), PC3, PC4 and motif speed (**Figure S5**). PC2 separated middle-stage MitoPark mice from the other groups, including Anxa1-tetanus mice (**Figure 5G**-**5I**).

To further understand the parameters that could explain how Anxa1+ DAN silencing resulted in a prodromal-like PD motor phenotype, we ranked the motifs according to their speed distribution and compared them across the different groups and their respective controls (**Figure 5J**). Although our recordings showed that Anxa1+ activity in dCP strongly correlated with action speed, we found that Anxa1-silenced mice did not show a significant reduction in all vigorous high-speed motifs. Specifically, we found that Anxa1-silenced mice displayed significant downregulation of five high-speed motifs (42, 34, 16, 0 and 25) (**Figure 5J**). When we compared the motif usage across the different mouse models, we found a subset of motifs that were significantly reduced across all the PD models (motifs 42 and 34), while another subset was reduced only in the middle-staged MitoPark group (motif 0 and 25; **Figure 5J**). Interestingly, for the actions defined as ‘rear’ and ‘turn’, the reduced motifs 42 and 25 represented the highest-speed motifs in that category, while for the ‘walk’ action category the significantly reduced motifs (motifs 0, 16, and 34) were not the highest-speed motifs within that category. This indicates that Anxa1+ DAN silencing did not reduce all high-speed or vigorous actions, suggesting that the reduction in locomotor speed observed in Anxa1-tetanus mice is not due to a general reduction of all high-speed motifs (**Figure 5J**). Additionally, the stationary-related motifs (3, 39, 27, and 28) were significantly increased only in the middle-stage MitoPark group, suggesting that the Anxa1+ DAN deficit does not influence the duration of stationary-associated behavior (i.e. akinesia/rigidity) (**Figure 5J**).

To better understand the role of Anxa1+ DANs in maintaining the action structure and a possible role in the prodromal motor deficits, we compared the transition probabilities between motifs. We did not observe any major effects in the overall structure of motif sequences after Anxa1+ DANs silencing (**Figure S5**); however, we identified that the significantly reduced high-speed “walk” motifs formed a small network with high-probability transition connections (motif 0 -> motif 16 -> motif 34; **Figure 5K**-**5M; Supplementary Video2**). Based on these results, we concluded that the Anxa1+ DANs are necessary for the expression of a subset of vigorous movements, and that their loss is associated with a specific bradykinetic profile that is behaviorally similar to the prodromal or early PD phenotype.

### Anxa1+ DANs are necessary for procedural motor learning

Loss of DA signaling in PD affects synaptic plasticity in the dorsal striatum, which may account for the motor learning deficits observed in PD patients^51^. Having established that Anxa1+ DANs do not encode motivational value, we next investigated their role in learning motivated behaviors. To this end, we first tested whether silencing Anxa1+ DANs would disrupt procedural learning in a simple 2-choice maze reversal task (**Figure 6A**). In this paradigm, water-restricted mice were placed in a modified T-maze with sharper angles at the choice points, referred to as the “arrow maze” (**Figure 6A**). The task required mice to initiate a trial and navigate to a designated arm, which consistently provided a water reward across days^52^ (**Figure 6A**). Similar to the mild motor effects during open field exploration, we found that Anxa1-silenced mice displayed only minor changes in locomotion, showing a small decrease in linear speed on the fourth day of the test without any changes in immobility time (**Figure 6B and** **6C; Figure S6**). Control mice successfully learned the basic task strategy over the four days, as evidenced by a significant increase in the preference for the rewarded arm and a significant increase in task efficiency quantified as the received reward relative to the traveled distance (**Figure 6B**-**6D; Figure S6**). However, mice with silenced Anxa1+ DANs failed to show any increase in the preference for the rewarded arm and did not improve their motor efficiency in the task (**Figure 6B**-**6D; Figure S6**). Our findings therefore establish that the activity of Anxa1+ DANs is essential for procedural learning when mice need to learn the basic task strategy to obtain rewards in a maze.

**Figure 6.**
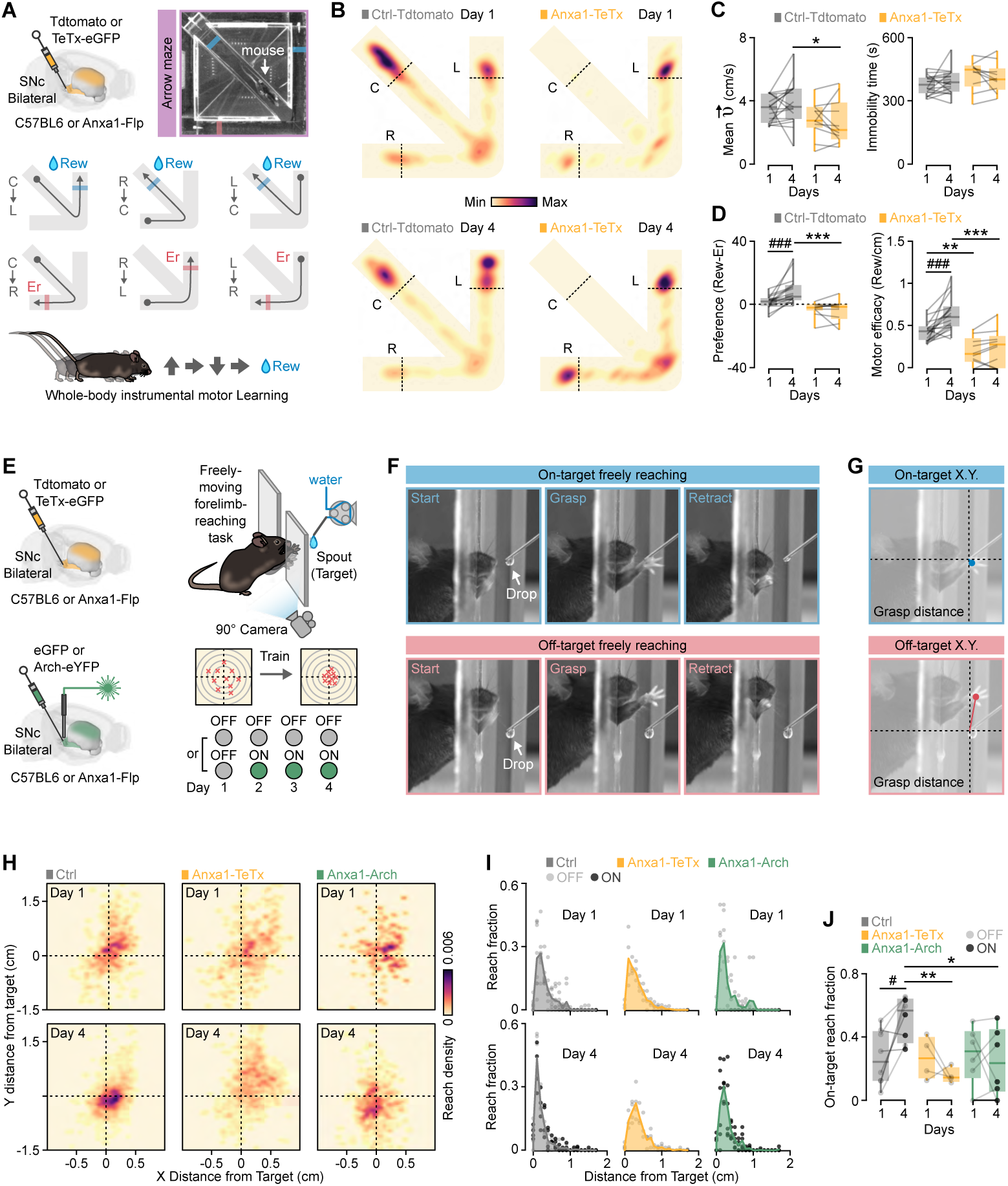
Anxa1+ DANs are required for procedural motor learning. **(A)** Illustration and image depicting the experimental setup for the 2-choice arrow maze task. **(B)** Representative heatmaps showing the position density of Ctrl-Tdtomato (left) and Anxa1-TeTx (right) mice on the first day (top) and fourth day (bottom) in the arrow maze task. **(C)** Boxplots displaying mean linear speed (left) and immobility time (right) for Ctrl-Tdtomato (n = 18) and Anxa1-TeTx (n = 10) mice on the first and fourth days of the arrow maze task. (RM two-way ANOVA for mean linear speed, Group effect: F(1,26) = 6.198, p = 0.0195; Ctrl-Tdtomato vs. Anxa1-TeTx: *p = 0.0473, Sidak’s post-hoc test). **(D)** Left: Boxplot illustrating the difference between rewarded and error transitions in Ctrl-Tdtomato (n = 18) and Anxa1-TeTx (n = 10) mice on the first and fourth days of the arrow maze task (RM two-way ANOVA, Group × Days: F(1,26) = 6.563, p = 0.0166; Day 1 vs. Day 2: ###p = 0.0009, Ctrl-Tdtomato vs. Anxa1-TeTx: **p < 0.0001, Sidak’s post-hoc test). Right: Boxplot illustrating locomotor efficacy for Ctrl-Tdtomato (n = 18) and Anxa1-TeTx (n = 10) mice on the first and fourth days of the arrow maze task (RM two-way ANOVA, Group × Days: F(1,26) = 6.065, p = 0.0207; Day 1 vs. Day 2: ###p < 0.0001, Ctrl-Tdtomato vs. Anxa1-TeTx: *p = 0.0027 and **p < 0.0001, Sidak’s post-hoc test). **(E)** Illustration representing the groups of mice (left) and the experimental design (right) for the forelimb-reaching task. **(F)** Example sequence of reaching efforts illustrating the three phases of movement: start, grasp, and retract, demonstrated in on-target (top) and off-target (bottom) reaching efforts. **(G)** Images depicting the method utilized to assess the performance of mice in the reaching task. **(H)** Heatmap showing the density of reaching efforts around the target for all groups of mice on the first (top) and fourth (bottom) days of the task. **(I)** Graph presenting the reach fraction distribution across 0.1 cm distance bins from the target for Ctrl mice (n = 8), Anxa1-TeTx mice (n = 4), and Anxa1-Arch mice (n = 6) on the first (top) and fourth (bottom) days of the task. **(J)** Boxplot summarizing the target accuracy fraction (defined as < 0.1 cm distance) for all groups of mice (RM two-way ANOVA, Group × Days: F(2,14) = 5.301, p = 0.0181; Day 1 vs. Day 2: #p = 0.0123, Ctrl vs. Anxa1-TeTx: **p = 0.0063, Ctrl vs. Anxa1-Arch: *p = 0.0268, Sidak’s post-hoc test). Boxplots display all data points, the 25th and 75th percentiles (box), the median (center), and the maxima (whiskers). Abbreviations: C: Central arm, L: Left arm, R: Right arm, DANs: Dopaminergic neurons, Ctrl: Control, TeTx: Tetanus toxin. Rew: reward, Er: error. See also **Figure S6** and **Supplementary Video3.**

To directly examine how Anxa1+ DANs shape the learning and execution of fine motor movements, we implemented a water drop reaching-grasping task^53^. In this task, water-restricted mice were trained to extend their forelimb through a small opening to reach a water droplet (**Figure 6E**). After a habituation period, mice underwent four training sessions (15 minutes each, one per day) to grasp the water droplet from a spout (**Figure 6E**). Anxa1+ DANs were specifically silenced using either the tetanus toxin light chain or the inhibitory opsin Arch in Anxa1-Flp mice (**Figure 6E**). Continuous optogenetic inhibition was applied to Anxa1-Arch mice during the last three training sessions (**Figure 6E**). Each reaching attempt began with the forelimb approaching the target (start phase), followed by digit extension, and grasping upon contact with the water droplet (grasp phase), and concluded with the retraction of the forelimb to consume the droplet (retract phase, **Figure 6F**; **Supplementary Video3**). Reaching accuracy was measured as the percentage of grasping attempts made within the target (< 0.2 cm) with the preferred forelimb (**Figure 6G; Figure S6**). As expected, control mice improved their reaching accuracy over the four training sessions (**Figure 6H**-**6J; Figure S6**). In contrast, Anxa1-tetanus mice exhibited normal accuracy during the first session but failed to refine their grasping movements in subsequent sessions (**Figure 6H**-**6J; Figure S6**). Comparison of motor learning impairments between genetic silencing and optogenetic inhibition revealed similar deficits, as Anxa1-Arch mice showed a significant reduction in reaching accuracy during the last three sessions (**Figure 6H- 6J; Figure S6**). These findings establish that Anxa1+ DAN activity is essential for procedural motor learning as a mechanism for improving dexterous motor skills.

## Discussion

It is widely accepted that SNc-DANs are the first to degenerate in PD^9^, yet the possible cell-type specific vulnerability remains a key question. In this study, we used an unbiased approach to identify the molecular signature of cell-type specific vulnerability, which revealed a specific subpopulation with expression of the marker Anxa1. The Anxa1 vulnerability profile was found across several mild or progressive PD models with different neurodegenerative mechanisms such as mitochondrial dysfunction^41,42^, catecholaminergic neurotoxin exposure^54^, and protein aggregation linked to Lewy body-like pathology^55^. RNA sequencing data from the early onset of degeneration in two progressive mouse PD models (MitoPark and conditional Mfn2 knockout mice) revealed a shared signature of significant Anxa1 downregulation. Importantly, we also found a specific and significant reduction in Anxa1 in iPSC-derived DANs from PD patients. Our functional investigation further established that Anxa1+ DANs do not encode reward-related signals and do not shape the motivation to work for natural rewards. Instead, the activity of Anxa1+ DAN axons was strongly correlated with high-speed and vigorous movements. Interestingly, genetic silencing of Anxa1+ DANs led to only moderate locomotor impairments, resembling a bradykinesia profile found after mild DAN degeneration as well as in MitoPark mice in the early degenerative stages. Behavioral segmentation and analysis of the organization of motif sequences revealed that silencing of Anxa1+ DANs resulted in significant reduction only in a specific subset of vigorous movement sequences—a pattern similarly found in the prodromal PD phase in mouse models. When we investigated the function of Anxa1+ DANs in cognitive and motor learning tasks, we found that they were necessary for procedural motor learning. Silencing Anxa1+ DANs disrupted the ability to learn the basic procedural strategy in a maze task and disrupted the refinement of reaching and grasping movements. Our findings therefore establish that Anxa1+ DANs are essential for learning and optimization of goal-directed motor behaviors.

Postmortem and neuroimaging studies in humans have pointed out that across SNc-DANs those located in nigrosome-1 of the ventral tier are the first to degenerate in PD^9,56^. A recent study showed that the non-human primate equivalent of nigrosomes contains Aldh1a1+ DANs^20^, and it is therefore likely that Anxa1 expression can define the vulnerable nigrosomes territories in primate SNc. Supporting this notion, the reported nigrosome-located DANs possess long SNr- penetrating dendrites with dorsoventral trajectories^20^, characteristic of the dendritic structures we found in Anxa1+ DANs. Interestingly, these dendritic formations are lost in a macaque PD model^20^, similar to the loss of Anxa1+ “dendrons” in our mild 6-OHDA model. In addition to their unique dendritic morphology, we found that Anxa1+ DANs targeted a specific dorsal domain in dCP spanning from the dorsomedial (DMS) to the dorsolateral (DLS) striatum, as reported previously^48^. This projection pattern corresponds to a possible vulnerable territory in CP, as the DA loss is most pronounced in the dorsal regions located just beneath the white matter in PD patients^57,58^. In summary, the neuroanatomical organization of Anxa1+ DANs defines a vulnerable nigrostriatal circuit in PD.

DA is in the reinforcement learning framework a reward-related teaching signal that can drive synaptic plasticity, to form action-value representations that guide motivated behavior. Midbrain DANs can encode reward prediction errors^59–61^, supporting the role of the DA system in reinforcement learning^62^. However, this theoretical framework does not account for the role of DA in motor or action initiation, behaviors that are impaired in PD patients^63^. Moreover, the idea that midbrain DANs transmit a scalar and homogeneous signal is continuously debated, and several studies show that DANs exhibit complex or non-reward related responses depending on their anatomical location^14,15,30,48,62^. We observed that Anxa1+ DANs showed a ramping activity signal that sharply declined directly after contact with the water droplet spout, which can be contrasted with the immediate and robust increase in the DA signal in central regions of CP (i.e. outside the Anxa1+ axon target area). Our findings agree with recordings in head-fixed mice, which show absence of reward-related signals in Anxa1+ striatal projections^48^, suggesting that the observed Anxa1+ DA ramping signals can signal reward approach in freely moving mice. In contrast to the lack of clear value or reward signals, we found that Anxa1+ DAN activity is strongly correlated with locomotor speed and vigorous actions during open field exploration. These findings align with previously described locomotion-reward gradient in the CP observed in head-fixed setups^30,48,64^, and we further validate these results in a freely-moving conditions. While functional evidence supports that stimulation of SNc DANs promotes rewarding and motivational effects^65–67^, we found that the selective optogenetic activation of Anxa1+ DANs could not reinforce motor actions. Our findings collectively establish that Anxa1+ DANs do not play a significant role in modulating reward-related or motivated behaviors through value signals.

Loss of SNc DANs projecting to the CP is associated with deficits in both movement initiation and invigoration, leading to akinesia/rigidity and bradykinesia, as observed in PD patients. A critical question is how the loss of Anxa1+ DANs specifically contributes to the PD behavioral phenotype. One of the key findings from our study is that silencing Anxa1+ DANs selectively suppresses a distinct subset of high-speed locomotor motifs, despite the strong correlation between Anxa1+ DAN activity and overall velocity. Notably, Anxa1-silenced mice retained the ability to perform high-speed motifs comparable to their control littermates, suggesting that these neurons do not directly regulate velocity. This phenotype mirrors PD-associated bradykinesia, which arises from an increased propensity for slow movements, as low-energy expenditure actions are preferentially reinforced, even though the individual retains the capacity to perform normal high-speed movements^3,34^. Another key finding of our study is that blocking DA release from Anxa1-terminals did not impact the motifs representing stationary behavior. Supporting this idea, the Anxa1+ axonal activity was not modulated at the onset of stationary motifs; instead, these motifs were associated with a significant increase in DA release in a striatal region outside the Anxa1+ axonal territory (i.e. cCP). In line with these observations, we found that stationary motifs become prominent in the more advanced stages of DAN degeneration, which extends beyond the loss of Anxa1+ DANs. These results are consistent with studies that have observed DAN calcium transients during deceleration^48^, as well as reduced SNc DAN activity at the initiation of movement^68^. We hypothesize that the stationary-related motifs represent akinesia/rigidity, motor symptoms that appear in advanced stages of degeneration^69^. In conclusion, we propose that the loss of Anxa1+ DANs results in an early bradykinetic phenotype characteristic of the prodromal PD, while akinesia/rigidity arises from the progressive degeneration of other SNc DANs in later disease stages.

DA-mediated synaptic plasticity in BG can act as a mechanism for procedural motor learning to reinforce and refine the association between specific motor plans, prediction signals, and action execution^51,70,71^. Striatal neurons demonstrate movement direction-selective tuning during both active and passive limb movements, underscoring their role in integrating sensory proprioceptive signals with actions^72,73^. DAN degeneration might impair this process, as PD patients show deficits in visuospatial processing^74^ and motor tasks^75^, for example precise grip-lift task^76^ indicating impairments in sensorimotor processing. The observed procedural motor learning deficits in PD patients^77^ suggest dysfunction in the perception of limb and body movements (i.e. kinaesthesia)^78,79^, potentially attributable to the loss of Anxa1+ DANs. Given that dCP neurons can encode the preparation and trajectory of body and limb movements including multisensory signals (visual and somatosensory)^80–83^, it is plausible that Anxa1+ DANs provide the error-based learning signals through their discrete axonal projections in dorsal striatal subregions. Our study suggests that Anxa1+ DAN function is essential for maintaining procedural motor learning and motor skills in prodromal PD, providing a targeted circuit approach to counteract the progressive worsening of severe motor deficits.

An interesting aspect for future investigation is the function of the Anxa1 protein, specifically considering the link between ANXA1 mutations and PD^84,85^. Anxa1 is a member of the annexin superfamily, characterized by its calcium-dependent binding to phospholipids^86^. Given that calcium influx during the pacemaking cycle of SNc DANs is a vulnerability factor compared to the resistant VTA neurons^87,88^, it is possible that the Anxa1-calcium interaction is a vulnerability mechanism in Anxa1+ SNc DANs – and preserving Anxa1+ DAN function can be a promising strategy to counteract the prodromal symptoms in PD.

## Methods

### Animals

All experimental procedures were approved by Stockholm’s Animal Experimentation Ethics Committee (Stockholms Norra Djurförsöksetiska Nämnd, approval numbers N166/15, 15440- 2020, and 21392-2021) and conducted in compliance with ethical guidelines. Mice used in this study were maintained on a C57BL/6N genetic background and housed in air-conditioned rooms at 20°C with 53% humidity, under a 12-hour light/dark cycle, with ad libitum access to food and water, except when on a water restriction schedule. Male and female mice aged 2-6 months were included in the experiments.

Anxa1-flp mice were generated by homologous recombination of a cassette with the flpo sequence in C57BL/6N ES cells (Cyagen). C57BL/6N mice were purchased from Charles River Laboratories. Mice homozygous for the loxP-flanked Tfam allele (Tfam^loxP/loxP^)^89^ were crossed with heterozygous DAT^IRESCre^ mice, resulting in double heterozygous offspring, which were then crossed with Tfam^loxP/loxP^ mice to produce MitoPark and control mice^42^. To generate mice with a conditional, full deletion of the gene Mfn2 in DANs (cMfn2KO), DATcreERT2^90^ mice were crossed with floxed Mfn2 (Mfn2^loxP/loxP^) mice^91^. In a subset of these crosses, the Gt(ROSA26)Sor^Stop–mito–YFP^ allele (stop-mitoYFP)^92^, encoding mitochondrially targeted YFP, was introduced at the parental generation. To generate mice with a conditional, full deletion of the autophagy gene Atg7 in DANs (cAtg7KO), DAT-Cre (DAT^IRESCre^) mice were crossed with floxed Atg7 (Atg7^loxP/loxP^) mice^43^.

### Stereotaxic surgery

Mice were anesthetized with isoflurane (4% induction and 1.5-2.5% maintenance) and placed in a stereotaxic frame (Harvard Apparatus, Holliston, MA). The analgesic Buprenorphine (0.1 mg/kg) was administered subcutaneously, and a local analgesic, Xylocaine/Lidocaine (4 mg/kg), was applied before the initial incision. A heating pad maintained the mice’s temperature at approximately 37°C, and eye ointment was applied to protect the eyes. For viral injections, a capillary attached to a Quintessential Stereotaxic Injector (Stoelting, Wood Dale, IL) was used. The capillary was held in place for 5 minutes post-injection before being slowly retracted from the brain. Carprofen (5 mg/kg) was administered at the end of the procedure as a postoperative analgesic, followed by a second dose 18-24 hours after surgery.

### Viral injections

Neuronal anatomical organization was assessed through bilateral or unilateral injections of 300 nL AAV1-Ef1a-fDIO-EYFP (Cat# AV55641, Addgene) into the SNc at coordinates AP −3.09, ML ±1.25, DV −4.2. For optogenetic experiments, 300 nL of AAV8-nEF-Coff/Fon-ChRmine-oScarlet (Cat# 137160, Addgene) was bilaterally injected into Anxa1-flp mice, while 300 nL of AAV8-nEF-Con/Foff 2.0-ChRmine-oScarlet (Cat# 137161, Addgene) was injected into DAT-Cre mice at the same coordinates. For FP recordings, unilateral injections of 300 nL or 500 nL AAV-DJ/2-hSyn1-chl-dFRT-jCaMP8m(rev)-dFRT-WPRE-bGHp (v740-DJ/22887, VVF Zurich) were administered to Anxa1-flp mice, while 300 nL of pGP-AAV9-CAG-FLEX-jGCaMP8m-WPRE (Cat# 162381, Addgene) was used in DAT-Cre mice. DA release recordings were conducted by injecting 500 nL of AAV5-CAG-dLight1.3b (Cat# 125560, Addgene) unilaterally into C57BL6 mice. For permanent inhibition of neurotransmission, 300 nL of ssAAV-9/2-hSyn1-chl-dFRT-TeTxLC_2A_NLS_dTomato(rev)-dFRT (v897-9, VVF Zurich) was bilaterally injected into Anxa1-Flp mice, with AAV5-CAG-TdTomato (Cat# 59462, Addgene) serving as the control. Short-term somatic inhibition was achieved by injecting 300 nL of AAV8-nEF-Coff/Fon-Arch3.3-p2a-EYFP (Cat# 137150, Addgene) into Anxa1-Flp mice, with AAV5-hsyn-GFP (Cat# 59462, Addgene) as control.

### Mild 6-OHDA lesion model

Mice were pretreated with desipramine (25 mg/kg, i.p.; Cat# D3900, Sigma–Aldrich) and pargyline (5 mg/kg, i.p.; Cat# P8013, Sigma–Aldrich) before being placed in a stereotaxic frame. A 1 μL injection of 6-OHDA (0.6 μg/μL in 0.02% ascorbate, Cat# 162957, Sigma–Aldrich) was administered into the medial forebrain bundle of the right hemisphere, using the coordinates AP −1.2, ML +1.25, DV −4.75 relative to bregma and the dural surface. Histological or behavioral assessments were conducted at least two weeks post-injection. Post-operative care included daily monitoring for one week, with sweetened milk (1:3 dilution in tap water) provided in the home cage to support recovery.

### Implant placement

Mice were implanted with either an optic fiber (200 μm, cerramic ferrule, NA 0.22, RWD, Cat# R-FOC-BL200C-22NA) or a fiber optic cannula (400 μm, cerramic ferrule, NA 0.22, Doric) after virus injection at the same stereotaxic surgery session. Optic fibers used in optogenetic experiments were positioned in SNc in coordinates: AP −3.09, ML +/- 1.25, DV −3.9. Fiber optic cannulas used in fiber photometry recordings were placed in CPu in coordinates: AP +0.74, ML+/−1.5, DV: −2.2. Following stereotaxic placement, the implants were fixed using Optibond primer and UV- curing liquid dental cement (Plurafill Flow+ A1, Cat# 246124). Mice were monitored to approximately one week post operation for recovery.

### Fluorescent in situ hybridization

Fluorescent in situ hybridization (ISH) using RNAscope was performed on 12 μm fresh frozen cryostat sections (Leica) with the RNAscope Multiplex Fluorescent Assay (ACD, Cat# 320850) or RNAscope Multiplex Fluorescent Assay v2 (ACD, Cat# 323110). Thaw-mounted sections were post-fixed in 4% paraformaldehyde for 15 minutes at 4 °C, then dehydrated in graded alcohol solutions. Protease IV (ACD, Cat# 322340) was applied for 30 minutes at room temperature, followed by probe hybridization for Anxa1 (Mm-Anxa1, Cat# 509291), Aldh1a1 (Mm-Aldh1a1-C2, Cat# 491321-C2), and Th (Mm-Th-C3, Cat# 317621-C3) for 2 hours at 40 °C. After hybridization, sections underwent four standardized amplification steps: Amp 1-FL for 30 minutes at 40 °C, Amp 2-FL for 15 minutes at 40 °C, Amp 3-FL for 30 minutes at 40 °C, and Amp 4C-FL for 15 minutes at 40 °C. Following the final amplification step, sections were counterstained with DAPI for 30 seconds and mounted with coverslips using Mowiol mounting medium. Imaging was performed on an LSM 880 confocal microscope (Carl Zeiss) using either a 20× 0.8 NA objective or a 63× 1.4 NA oil immersion objective. Z-stacks with a thickness of 6–9 μm were captured for each imaged area.

### Immunofluorescence

Perfused brains (4% paraformaldehyde) were sectioned at 50-60 µm using a vibratome (Leica VT1000, Leica Microsystems, Nussloch GmbH, Germany). Sections were washed twice for 5 minutes each in phosphate-buffered saline (PBS) and then incubated in blocking buffer (10% donkey serum, 0.3% Triton-X in PBS) for 1 hour. Following this blocking step, the sections were incubated overnight at 4°C with primary antibodies: rabbit anti-Anxa1 (Cat# 71-3400, Thermo-Fisher), goat anti-Aldh1a1 (Cat# AF5869, R&D Systems), rabbit anti-MOR (Cat# ab17934, Abcam), chicken anti-TH (Cat# ab76442, Abcam), mouse anti-TH (Cat# AB152, Sigma-Aldrich), rabbit anti-GFP (Cat# A6455, Invitrogen), chicken anti-GFP (Cat# A10262, Invitrogen), mouse anti-p62 (Cat# 610833, BD Biosciences) and rabbit anti-RFP (Cat# 600-401-379-RTU, Rockland), all diluted in 2.5% donkey serum and 0.3% Triton-X in PBS. The next day, sections were washed three times for 5 minutes each in PBS and incubated with fluorescent-conjugated secondary antibodies (anti-rabbit, anti-mouse, anti-chicken, or anti-goat) for 2 hours. Following three additional 5-minute washes in PBS, sections were mounted on coated slides and covered with coverslips using Mowiol mounting medium. Imaging was performed using either an LSM 880 confocal microscope (Carl Zeiss) with a 20× 0.8 NA objective or a Leica DM6000 B microscope (Leica Microsystems) with a 10× objective. For confocal imaging, Z-stacks of 15–20 µm thickness were acquired for each captured region.

### Anatomical tracing

Confocal microscope images were stitched using Zeiss software, followed by processing in ImageJ for brightness and contrast adjustment. Cell counting was performed with Imaris software. X-Y coordinates of distinct neuronal populations were utilized to generate 2D graphical representations of the anatomical organization within the SNc. 2D representations of dendrites and axonal distributions were created in ImageJ as previously described^13^. 3D graphical reconstructions of axons and dendrites distributions were created using cell registration software in Napari, developed in-house.

### Isolation of DANs for bulk RNAseq

Brains were dissected from mitoYFP expressing mice, sectioned and dissociated into single cell suspensions, as previously described^93^. After dissociation, mitoYFP positive and negative cells were collected using a BD FACSAria III Cell Sorter for RNAseq analysis.

### RT-qPCR of hiPSC-DANs

iPSC-derived DANs were generated from healthy human donors or Parkinson’s patients with SNCA locus triplication, following an expanded version of Krik’s protocol as previously described^94^ adapted from Kriks et al.^95^ and Fedele et al.^96^. Briefly, iPSCs were differentiated into ventral midbrain neural precursor cells, expanded, and matured to achieve DAN identity. On Day 70, cell lysates were collected using ice-cold PBS, and mRNA was extracted with the Qiagen RNeasy RNA purification kit (Cat# 74104) per manufacturer instructions. RNA quantification was performed using a DS-11 Nanodrop, and cDNA synthesis was conducted using the Invitrogen SuperScript III One-Step RT-PCR System kit (Cat# 12574026). RT-qPCR was performed on an Applied Biosystems StepOnePlus Real-Time PCR system with Fast SYBR Green Master Mix (Cat# 4385612), and mRNA levels were normalized to the housekeeping gene B2M as described in Matsuzaki et al.^97^.

### Open field test

Mice were placed in a 40×40 box with a transparent floor and dark walls. Mouse locomotion was recorded from below with an Oryx 10GigE camera (30 fps, 800 x 800 px; Hamamatsu Photonics). All lights in the room were dimmed, and the arena was illuminated by a white LED light strip glued along the rim of the clear floor.

### Fixed ratio & Progressive ratio protocol

Water-restricted mice were placed in a 10 × 10 cm box equipped with a single port containing a waterspout under dim light conditions. Mice performed nosepokes to receive a reward of 1 water droplet (2 µl) over a 15-minute session. The reward schedule began with a fixed ratio, where 1 nosepoke resulted in 1 reward on the first day, 2 nosepokes per reward on the second day, and 3 nosepokes per reward on the third day. On the fourth and fifth days, a fixed ratio of 5 was implemented, requiring 5 nosepokes for each reward. During the final two days, a progressive ratio protocol was introduced, with the response requirement for each reward following a doubling pattern, where each entry requirement repeated once before doubling. This schedule was defined by the formula:

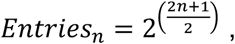

resulting in the following sequence: 1-2-2-4-4-8-8-16-16-32-32-64-64. Nosepoke counts were recorded using a custom-built counter that detected beam breaks at the port entrance.

### *In vivo* optogenetic manipulation

A Cobolt light source was connected to a low-autofluorescence patch cord (Doric, 200 μm) via ceramic sleeves (Thorlabs) to the optic fiber stubs implanted in the mouse. Laser power was adjusted by manually dialing an analog knob on the power supply. Before each experiment, light output at the tip of the ferrule in the patch cord was calibrated to the desired power using an optical power and energy meter and a photodiode power sensor (Thorlabs). Photostimulation frequency and duration were controlled using a custom-written Arduino script (Arduino IDE) managed through Bonsai (v2.6.3)^98^. In ChRmine experiments, optostimulation was delivered as 1-second pulses of red light (640 nm, 40 Hz, 5 mW). In Arch experiments, optoinhibition was applied as continuous green light (532 nm, 5 mW).

### Optogenetic self-stimulation

Mice were placed in a 10 × 10 cm box and allowed to freely perform nosepokes in two ports during a 15-minute session. On the first two days, nosepokes to the right port were paired with bilateral laser stimulation in the SNc (5 mW, 1 s, 40 Hz pulse train). For the following four days, laser stimulation was instead paired with nosepokes to the left port. Nosepoke counts were recorded using a custom-built counter that detected beam breaks at each port entrance.

### Arrow maze choice task

Water restricted mice were introduced to a modified T-maze consisting of a 40×40 cm box with a transparent floor, featuring two lateral corridors branching from a central one. Each corridor was equipped with a waterspout capable of delivering a 3 μL drop of water. At the initiation of every trial, the mice received one drop of water in the central corridor, after which they were required to choose between the two lateral corridors, with only one providing a reward. Over the course of four consecutive days, the mice were trained to obtain the reward from the left corridor of the maze. An LED light strip illuminated the arena. All sessions lasted 10 minutes. The task was implemented using the Bonsai software 43, used to detect entries into the corridor end zones from a live video feed, and two Arduino Micro microcontrollers (Arduino), controlling the waterspouts and communicating with Bonsai via the firmata protocol 44, respectively. Video frames were acquired at 30 fps using an Oryx 10GigE camera (Hamamatsu Photonics) placed below the arena

### Forelimb reaching task

Water-restricted mice were placed in a 10 × 10 cm enclosure with transparent walls. A 1-cm-wide opening on one wall provided access to a waterspout, dispensing water at intervals of 5-7 seconds. The training protocol began with 3 days of habituation, during which the waterspout was positioned directly at the wall (0 cm) and 5.5 cm from the floor. Following habituation, the water spout was gradually shifted 1 cm away from the wall, maintaining the same height, forcing mice to use their forelimbs to reach the water reward. Once mice consistently used their forelimbs to retrieve water, formal training commenced, continuing for 4 consecutive days in 15-minute sessions. For Arch experiments, the laser remained off during the initial 4 days (habituation and the first day of reaching) and was activated during the final 3 days of the experiment. Mice behavior was recorded daily using a FLIX camera (60 fps, 800 × 800 px) positioned laterally. Behavioral events and distances from the waterspout were manually analyzed and measured using Kinovea (v2023.1.2).

### Fiber photometry recordings

The used fiber photometry setup has been previously described in detail^99^. Briefly, calcium- or DA-dependent fluorescence signals were acquired at 60 frames per second (fps) using an excitation wavelength of 470 nm, alongside an isosbestic control signal at 405 nm. The light power of the 470 nm LED was between 20-30 μW, with the power of the 405 nm LED ranging slightly lower, usually between 10-25 μW. Signals from both wavelengths were normalized, filtered, and subsequently subtracted using DoricStudio software (V5). Artifacts and instances of signal intensity loss were excluded from the final data set. The resulting z-scored fluorescence signals were then aligned to specified behavioral events.

### Behavioral analysis

Open field, arrow maze and reaching behavioral tasks were implemented through Bonsai software 43^98^, which facilitated synchronization between behavioral videos and fiber photometry setup and/or laser driver (optogenetic experiments). Deep Lab Cut (DLC v2.3.7)^49^ software was used for tracking the mouse body parts: snout, four paws and the tail base, thus extracting the basic kinematic characteristics of the mouse: distance travelled, linear and angular speed and acceleration. For behavioral motif identification, the unsupervised deep learning framework VAME was trained on 195 videos (more than 1’500’000 frames) with a test fraction of 0.1, a time window of 30 frames, and a latent dimensionality of 30, converging at 50 epochs. Clustering analysis with 100 discrete states and a 1% motif usage threshold identified 47 relevant behavioral motifs in the dataset. Among these, 10 were merged with others due to their close similarity and minimal usage, resulting in a final set of 37 motifs. In the reaching test, behavioral events and distances from waterspout were manually identified and measured using Kinovea (v2023.1.2).

### Statistical analysis and graphs

Statistical analysis was performed by using two-sided unpaired t-test, two-sided paired t-test, one-sample two-tailed t-test, repeated measurements (RM) one-way analysis of variance (ANOVA) test, Kruskal-Wallis test, Friedman test, two-way ANOVA and RM two-way ANOVA using GraphPad Prism 10. P-value in multiple two-sided unpaired t-tests and one-sample two-tailed t-tests were adjusted using Holm-Sidak method (α=0.05) and Bonferroni correction respectively. Kruskal-Wallis test and Friedman were followed by Dunn’s post-hoc test. RM one-way ANOVA, two-way ANOVA and RM two-way ANOVA were followed by Sidak’s post-hoc test. Graphs were made in GraphPad Prism and R. All the illustrations shown in the Figure were made by using Inkscape 1.3.2. The 3D brain illustrations presented in the Figure were created and obtained from the Scalable Brain Atlas.

## Authors contributions

Conceptualization, K.M. and I.M.; Methodology, I.M., A.C., R.F.; Investigation, I.M., A.C., V.S., C.L, I.A.S., K.C., R.F.; Writing – Original Draft, I.M. and K.M.; Writing – Review & Editing, I.M., A.C., V.S., K.M.; Visualization, I.M., A.C., V.S.; Resources, R. W-M, P.M.; Supervision, K.M., R. W-M, P.M.; Funding Acquisition, K.M., R.F., R. W-M, P.M.; Project Administration: K.M.

## Acknowledgements

This research was funded in part by Aligning Science Across Parkinson’s ASAP-020370 through the Michael J. Fox Foundation for Parkinson’s Research (MJFF), the Swedish research council (grant 2018-00608 to K.M.), the Swedish Brain Foundation (Hjärnfonden grant to K.M.). I.M. was supported by the Swedish Society for Medical Research (815200-8317). A.C. was supported by the Swiss National Foundation (P500PB_214355) and the Amicitia Foundation. V.S. was supported by Karolinska Institutet doctoral funding (KID). K.M.L.C was supported by a Junior Research Fellowship from St. John’s College. R.F. was supported by the Swedish research council (grant 2022-01477), Hjärnfonden, Åhlén-stiftelsen, StratNeuro, and Hedlunds stiftelse. The authors thank Viktor Jonsson and Markus Ringnér, part of the National Bioinformatics Infrastructure Sweden at SciLifeLab and supported by the Knut och Alice Wallenberg Stiftelse, for technical assistance with RNAseq data analysis. The authors thank Alexander Wolthon for animal care and mouse colony management. For the purpose of open access, the authors have applied for a CC BY public copyright license to all authors accepted manuscript arising from submission.

## Code and data availability

The data, code, protocols, and key lab materials used and generated in this study are listed in a Key Resource Table alongside their persistent identifiers at Zenodo (http://doi.org/10.5281/zenodo.14138011). The RNA-seq dataset generated and analyzed during the current study will be available upon publication (NCBI Gene Expression Omnibus, accession number GSE282753).

## Declaration of generative AI and AI-assisted technologies in the writing process

During the preparation of the text, the authors used ChatGPT-4.0 for language and grammar corrections. After using this tool, the authors reviewed and edited the content as needed and take full responsibility for the content of the publication.

