## Supplementary Figures for "Anxa1+ dopamine neuron vulnerability defines prodromal Parkinson’s disease bradykinesia and procedural motor learning impairment"

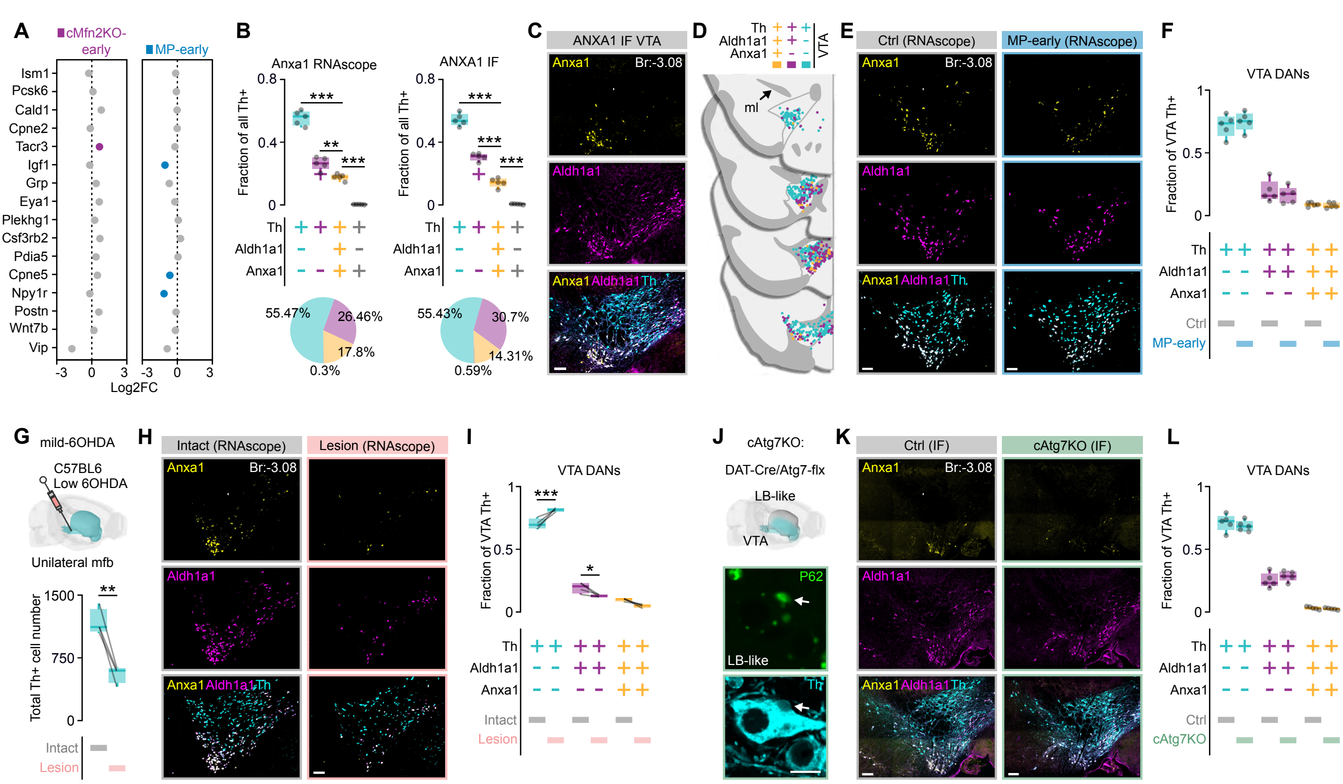

A
cMfn2KO-early
MP-early
Ism1
Pcsk6
Cald1
Cpne2
Tacr3
Igf1
Grp
Eya1
Plekhg1
Csf3rb2
Pdia5
Cpne5
Npy1r
Postn
Wnt7b
Vip
Log2FC
B
Anxa1 RNAscope
ANXA1 IF
Fraction of all Th+
Th
Aldh1a1
Anxa1
55.47%
26.46%
17.8%
0.3%
55.43%
30.7%
14.31%
0.59%
C
ANXA1 IF VTA
Anxa1
Br:-3.08
Aldh1a1
Anxa1 Aldh1a1 Th
D
Th
Aldh1a1
Anxa1
VTA
ml
E
Ctrl (RNAscope)
MP-early (RNAscope)
F
VTA DANs
Fraction of VTA Th+
Ctrl
MP-early
G
mild-6OHDA
C57BL6
Low 6OHDA
Unilateral mfb
Total Th+ cell number
Intact
Lesion
H
Intact (RNAscope)
Lesion (RNAscope)
I
VTA DANs
J
cAtg7KO:
DAT-Cre/Atg7-flx
LB-like
VTA
P62
Th
K
Ctrl (IF)
cAtg7KO (IF)
L
VTA DANs
cAtg7KO

**Figure S1. Anxa1 expression in VTA DANs, Related to Figure 1.**

**(A)** Graph depicting significantly downregulated & upregulated additional DAN subtype markers (blue for MP-early and magenta for cMfn2KO-early, FDR < 0.05).

**(B)** Boxplot showing the proportions of the three DAN subpopulations in the midbrain of Ctrl mice using ISH (right, n=5 mice; one-way ANOVA, F (3, 16) = 254.9, p < 0.0001; **p = 0.0036, ***p < 0.0001, Sidak's post-hoc test ) and IF (left, n=5 mice; one-way ANOVA, F (3, 16) = 364.7, p < 0.0001; ***p < 0.0001, Sidak's post-hoc test).

**(C)** IF images (left) and schematic representation (right) showing co-expression of Anxa1, Aldh1a1, and Th in the VTA (scale bars: 200 μm).

**(D)** Coronal sections illustrating the spatial distribution of these subpopulations in the VTA.

**(E)** ISH showing the expression of Anxa1, Aldh1a1, and Th in the VTA of Ctrl and MP-early mice.

**(F)** Boxplot showing proportions of the three VTA-DAN subpopulations on Ctrl (n=5 mice) and MP-early (n=5 mice) mice.

**(G)** Schematic representation (top) of the low-concentration 6-OHDA model and boxplot (bottom) showing the number of midbrain Th+ cells in intact and lesion sides (**p = 0.0039, Paired t-test).

**(H)** Representative images showing the expression of Anxa1, Aldh1a1, and Th in the VTA of mild 6-OHDA mice.

**(I)** Boxplot comparing the proportions of the three DAN subpopulations in the intact versus lesioned hemispheres (n = 4; RM two-way ANOVA, Side × Subtype: F(2,9) = 24.78, p = 0.0002; *p = 0.0263, ***p = 0.0009, Sidak's post-hoc test)

**(J)** Schematic illustration of cAtg7KO mice and IF images showing LB-like p62+ inclusions (arrows) in VTA DANs.

**(K)** IF images showing the expression of Anxa1, Aldh1a1, and Th in the VTA of Ctrl and cAtg7KO mice.

**(L)** Boxplot showing the proportions of the three VTA DAN subpopulations in Ctrl (n=5 mice) and cAtg7KO (n=5 mice) mice.

Boxplots display all data points, with the 25th and 75th percentiles (box), the median (center), and the maxima (whiskers).

Abbreviations: DANs: Dopaminergic Neurons, MP: MitoPark, cMfn2KO: Conditional Mitofusin 2 Knockout, FDR: False Discovery Rate, FC: fold change, VTA: ventral tegmental area, IF: Immunofluorescence, ISH: In Situ Hybridization, Ctrl: Control, 6-OHDA: 6-Hydroxydopamine, mfb: medial forebrain bundle, cAtg7KO: Conditional Atg7 Knockout, LB: Lewy Bodies.

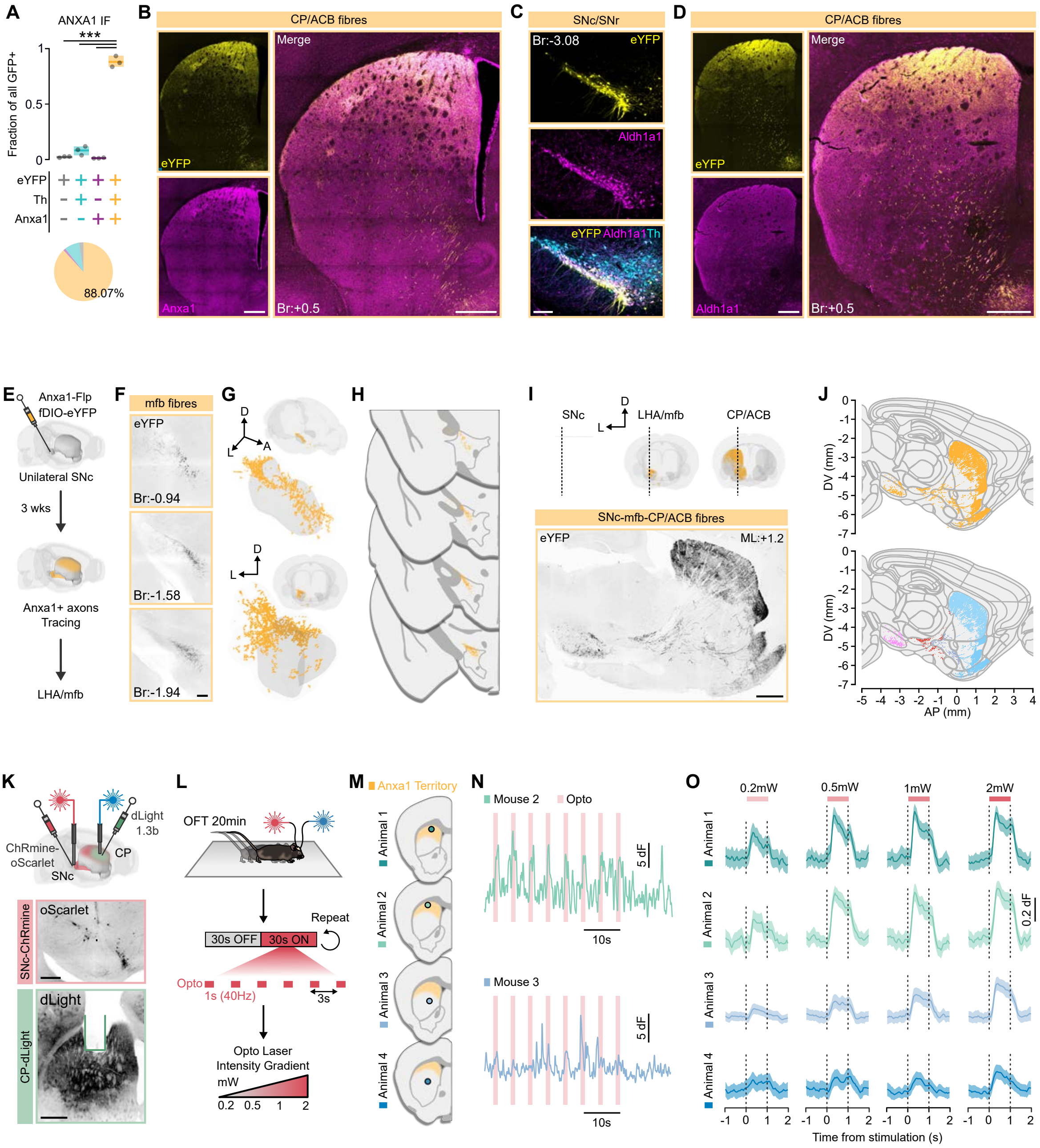
A
ANXA1 IF
***
Fraction of all GFP+
eYFP
Th
Anxa1
88.07%
B
CP/ACB fibres
eYFP
Anxa1
Merge
Br:+0.5
C
SNc/SNr
Br:-3.08
eYFP
Aldh1a1
eYFP Aldh1a1 Th
D
CP/ACB fibres
eYFP
Aldh1a1
Merge
Br:+0.5
E
Anxa1-Flp
fDIO-eYFP
Unilateral SNc
3 wks
Anxa1+ axons
Tracing
LHA/mfb
F
mfb fibres
eYFP
Br:-0.94
Br:-1.58
Br:-1.94
G
H
I
SNc
LHA/mfb
CP/ACB
SNc-mfb-CP/ACB fibres
eYFP
ML:+1.2
J
DV (mm)
AP (mm)
K
dLight
1.3b
ChRmine-
oScarlet
CP
SNc
SNc-ChRmine
oScarlet
CP-dLight
dLight
L
OFT 20min
Repeat
30s OFF
30s ON
Opto
1s (40Hz)
3s
Opto Laser
Intensity Gradient
mW
0.2 0.5 1 2
M
Anxa1 Territory
Animal 1
Animal 2
Animal 3
Animal 4
N
Mouse 2
Opto
5 dF
10s
Mouse 3
5 dF
10s
O
0.2mW
0.5mW
1mW
2mW
Animal 1
Animal 2
Animal 3
Animal 4
0.2 dF
Time from stimulation (s)

**Figure S2. Anatomical tracing of Anxa1+ projections across the brain, Related to Figure 2.**
**(A)** Graph showing the proportion of eYFP-labeled cells overlapping with Th and Anxa1 immunofluorescence (one-way ANOVA, $F(3, 8) = 517.4$, $p < 0.0001$; ***$p < 0.0001$, Sidak's post-hoc test).
**(B)** Fluorescent images of eYFP+ labeled fibers overlapping with Anxa1-labeled fibers in CP/ACB (scale bars: 500 µm).
**(C)** Immunofluorescence (IF) images showing eYFP labeling overlapping with Aldh1a1+ DANs (scale bar: 200 µm).
**(D)** Fluorescent images of eYFP+ labeled fibers overlapping with Aldh1a1+ labeled fibers in CP/ACB (scale bars: 500 µm).
**(E)** Schematic representation showing unilateral Anxa1+ fiber labeling.
**(F)** Fluorescent images showing eYFP+ labeled axons at different anteroposterior coronal levels of the LHA/mfb (scale bars: 200 µm).
**(G)** 3D reconstruction of eYFP+ labeled fibers within the LHA/mfb.
**(H)** Schematic coronal sections depicting the distribution of eYFP+ labeled fibers in the LHA/mfb.
**(I)** Fluorescent sagittal image showing the eYFP+ fiber labeling in SN, LHA/mfb and CP/ACB (scale bar: 500 µm).
**(J)** Graph showing the distribution of eYFP+ fibers in sagittal section (+1.2 mm lateral from midline) color-coded according to Allen Brain atlas region color.
**(K)** Illustration (upper) and fluorescent images (lower) showing the distinct groups of mice used for FP recordings in CP coupled with optogenetic stimulation (scale bars: 30 µm and 500 µm).
**(L)** Illustration (upper) showing the experimental setup of the photometry recordings coupled with optogenetic stimulation in an open arena. Optogenetic stimulation protocol (middle) and relative light intensity (lower) used in the experiment.
**(M)** Schematic coronal sections illustrating the recording locations in the CP for each mouse (n=4).
**(N)** Example traces showing FP recorded signal (dLight) during optogenetic stimulation.
**(O)** Z-score of the FP signals (dLight) aligned on optogenetic stimulation onset for each mouse in the experiment with distinct light intensity.
Abbreviations: IF: Immunofluorescence, Br: brema, eYFP: enhanced Yellow Fluorescent Protein, CP: Caudoputamen, ACB: Accumbens, SNc: substancia nigra pars compacta, SNr: substancia nigra pars reticulata, LHA: lateral hypothalamus, mfb: medial forebrain bundle, A: anterior, D: dorsal, L: lateral, ML: mediolateral, DV: dorsoventral, AP: anteroposterior, OFT: open field test, Opto: optogenetic stimulation.

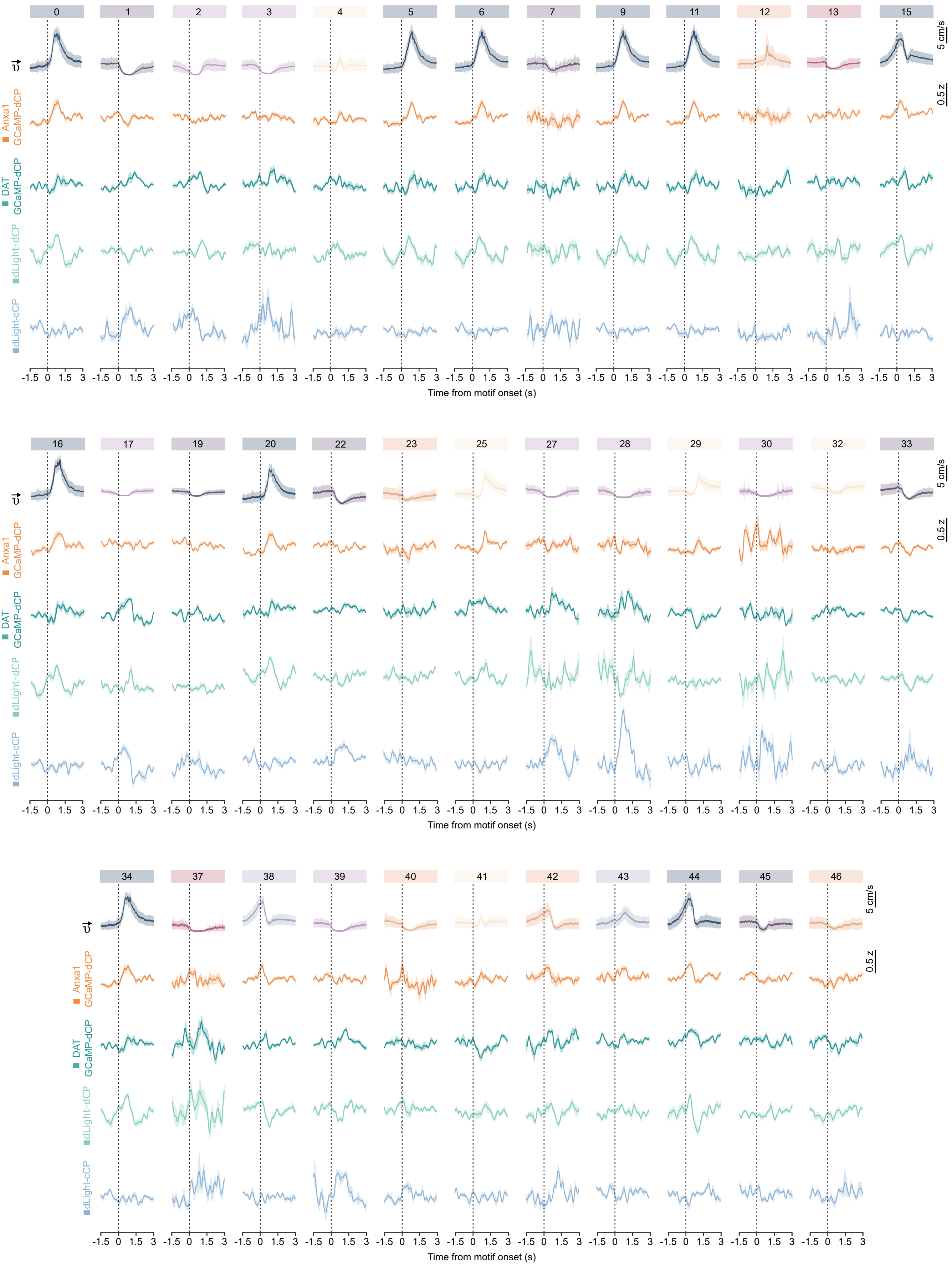

0 1 2 3 4 5 6 7 9 11 12 13 15
16 17 19 20 22 23 25 27 28 29 30 32 33
34 37 38 39 40 41 42 43 44 45 46
5 cm/s
0.5 z
Anxa1 GCaMP-dCP
DAT GCaMP-dCP
dLight-dCP
dLight-cCP
-1.5 0 1.5 3
Time from motif onset (s)

**Figure S3. Linear speed and FP signal traces aligned to the onset of different motifs, Related to Figure 4.**

Graph showing for each behavioral motif the linear velocity of the mice aligned with the z-score of the FP signals for each mouse group (Anxa1 GCaMP-dCP, DAT GCaMP-dCP, dLight-dCP and dLight-cCP).

Abbreviations: DAT: dopamine transporter, dCP: dorsal caudoputamen, cCP: central caudoputamen.

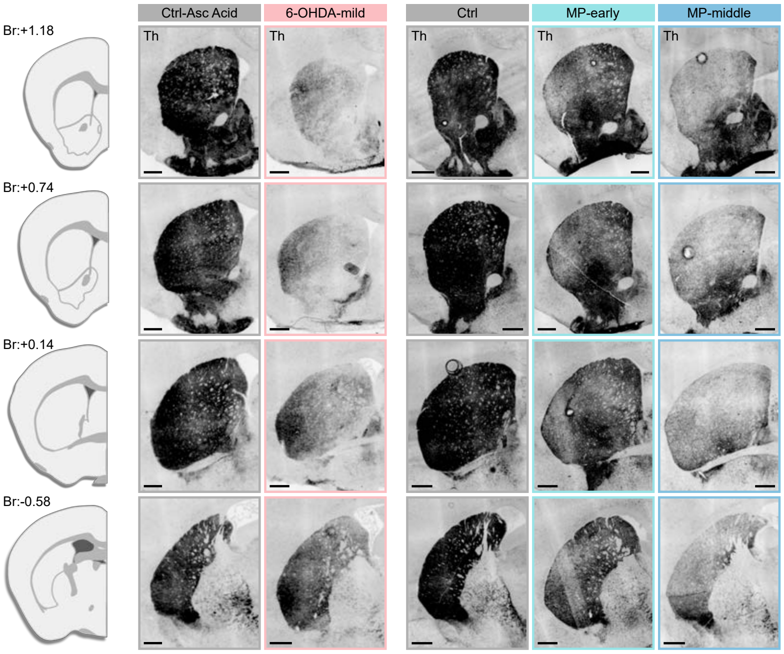

Ctrl-Asc Acid
6-OHDA-mild
Ctrl
MP-early
MP-middle
Br:+1.18
Br:+0.74
Br:+0.14
Br:-0.58
Th

**Figure S4. Tyrosine hydroxylase distribution in the CP/ACB of PD mouse models, Related to Figure 5.**

Immunofluorescence images showing the tyrosine hydroxylase (Th) expression in four anteroposterior levels of the striatum for the PD mouse models used in open field test: Ctrl Asc. Acid, 6-OHDA-mid, Ctrl, MP-early, MP-middle (scale bars: 500 μm).

Abbreviations: Br: bregma, 6-OHDA: 6-Hydroxydopamine, MP: Mitopark.

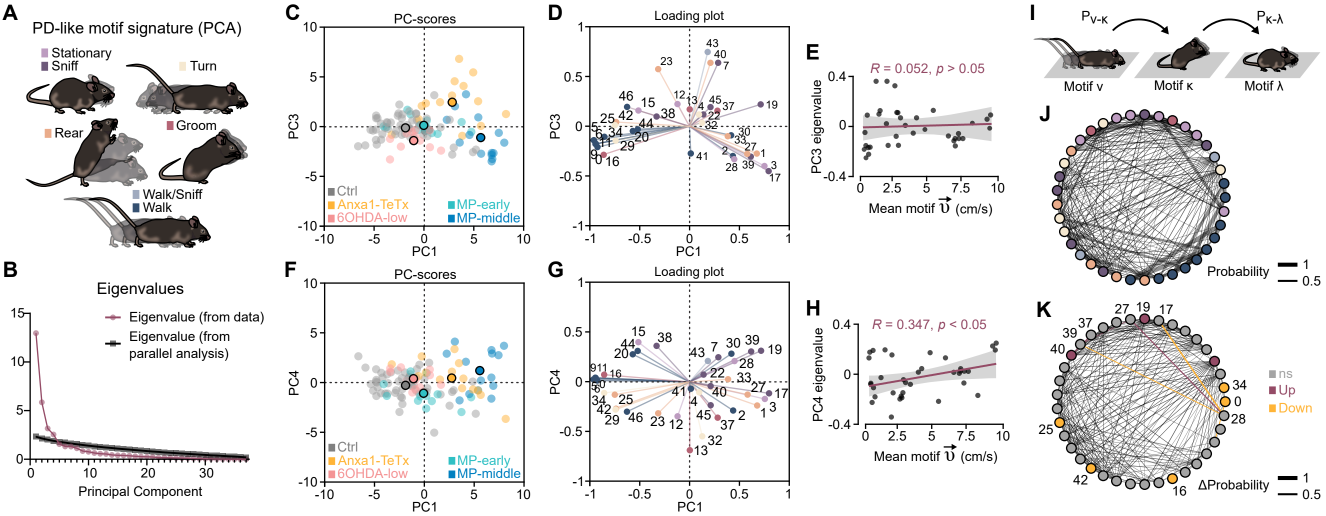
A
PD-like motif signature (PCA)
Stationary
Sniff
Turn
Rear
Groom
Walk/Sniff
Walk
B
Eigenvalues
Eigenvalue (from data)
Eigenvalue (from parallel analysis)
Principal Component
C
PC-scores
PC3
PC1
Ctrl
Anxa1-TeTx
6OHDA-low
MP-early
MP-middle
D
Loading plot
PC3
PC1
E
R = 0.052, p > 0.05
PC3 eigenvalue
Mean motif 𝜐⃗ (cm/s)
F
PC-scores
PC4
PC1
Ctrl
Anxa1-TeTx
6OHDA-low
MP-early
MP-middle
G
Loading plot
PC4
PC1
H
R = 0.347, p < 0.05
PC4 eigenvalue
Mean motif 𝜐⃗ (cm/s)
I
Pν-κ
Pκ-λ
Motif ν
Motif κ
Motif λ
J
Probability
K
ns
Up
Down
ΔProbability

**Figure S5. PCA and transition probability network analysis of motifs, Related to Figure 5.**
**(A)** Illustration showing the behavioral motifs groups extracted using VAME.
**(B)** Graph showing the eigenvalues of the principal components from the data and the parallel analysis.
**(C)** Principal component analysis based on motif usage reporting PC1 and PC3 for each group of mice and their controls.
**(D)** Loading plot showing the PC1 and PC3 for all the behavioral motifs.
**(E)** Principal component analysis based on motif usage reporting PC1 and PC4 for each group of mice and their controls.
**(F)** Loading plot showing the PC1 and PC4 for all the behavioral motifs.
**(G)** Graph showing the correlation of PC3 eigenvalues and the mean velocity of the behavioral motifs ($R = 0.052$, $p > 0.05$).
**(H)** Graph showing the correlation of PC4 eigenvalues and the mean velocity of the behavioral motifs ($R = 0.347$, $p < 0.05$).
**(I)** Illustration showing the probability of one motif occurring after the other.
**(J)** Statemap depiction of motifs (as nodes) and transition probabilities (as weighted edges from control. Statemap depiction of the difference between the control statemap and the Anxa1-tetanus statemap (n = 3 mice; red: significant upregulated transition, yellow: significant downregulated transition; Fisher's Exact Test, adjusted $p < 0.05$, Benjamini-Hochberg correction). The colored nodes represent the motifs with statistically significant altered usage in Anxa1-TeTx mice according to Figure 5 (red: significant upregulated motif, yellow: significant downregulated transition motif; multiple two-sided unpaired t-tests, adjusted $p < 0.05$, Holm-Sidak method).
Abbreviations: PD: Parkinson disease, Asc: ascorbic, PCA: principal component analysis, PC: principal component, R: Pearson correlation coefficient, VAME: Variational Animal Motion Embedding.

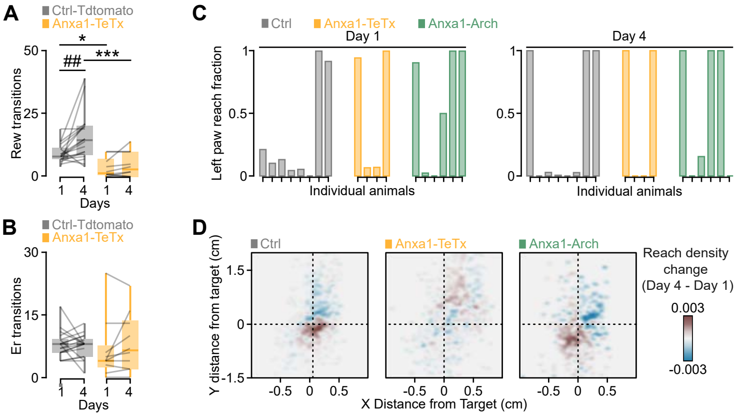

A
Ctrl-Tdtomato
Anxa1-TeTx
Rew transitions
Days
B
Er transitions
C
Ctrl
Anxa1-TeTx
Anxa1-Arch
Day 1
Day 4
Left paw reach fraction
Individual animals
D
Y distance from target (cm)
X Distance from Target (cm)
Reach density change (Day 4 - Day 1)
0.003
-0.003

**Figure S6. The role of Anxa1+ DANs in the maze and forelimb reaching task, Related to Figure 6.**

**(A-B)** Boxplots illustrating the number of rewarded transitions in Ctrl-Tdtomato (n = 18) and Anxa1-TeTx (n = 10) mice on the first and fourth days of the arrow maze task (RM two-way ANOVA, Group's effect: $F(1,26) = 17$, $p = 0.0003$, Day's effect: $F(1,26) = 6.554$, $p = 0.0166$; Day 1 vs. Day 2: ##p = 0.0025, Ctrl-Tdtomato vs. Anxa1-TeTx: *p = 0.0262, ***p=0.0001 Sidak's post-hoc test).

**(B)** Boxplots illustrating the number of error transitions in Ctrl-Tdtomato (n = 18) and Anxa1-TeTx (n = 10) mice on the first and fourth days of the arrow maze task.

**(C)** Bar plots showing the fraction of left forelimb reaching efforts per individual mouse.

**(D)** Heatmap showing the density change of reaching efforts around the target for all groups of mice between the first and fourth day of the task (red: increase in reach density on day 4, blue: decrease in reach density on day 4).

Abbreviations: Ctrl: control, Tetx: Tetanus toxin, Rew: rewarded, Er: error.
